# Distance-based metrics for comparing conformational ensembles of intrinsically disordered proteins

**DOI:** 10.1101/2020.04.06.027979

**Authors:** Tamas Lazar, Mainak Guharoy, Wim Vranken, Sarah Rauscher, Shoshana J. Wodak, Peter Tompa

## Abstract

Intrinsically disordered proteins (IDPs) are proteins whose native functional states represent ensembles of highly diverse conformations. Such ensembles are a challenge for quantitative structure comparisons as their conformational diversity precludes optimal superimposition of the atomic coordinates, necessary for deriving common similarity measures such as the root-mean-square deviation (RMSD) of these coordinates. Here we introduce superimposition-free metrics, which are based on computing matrices of Cα-Cα distance distributions within ensembles and comparing these matrices between ensembles. Differences between two matrices yield information on the similarity between specific regions of the polypeptide, whereas the global structural similarity is captured by the ens_dRMS, defined as the root-mean-square difference between the medians of the Cα-Cαdistance distributions of two ensembles. Together, our metrics enable rigorous investigations of structure-function relationships in conformational ensembles of IDPs derived using experimental restraints or by molecular simulations, and for proteins containing both structured and disordered regions.

**Statement of Significance:** Important biological insight is obtained from comparing the high-resolution structures of proteins. Such comparisons commonly involve superimposing two protein structures and computing the residual root-mean-square deviation of the atomic positions. This approach cannot be applied to intrinsically disordered proteins (IDPs) because IDPs do not adopt well-defined 3D structures, rather, their native functional state is defined by ensembles of heterogeneous conformations that cannot be meaningfully superimposed. We report new measures that quantify the local and global similarity between different conformational ensembles by evaluating differences between the distributions of residue-residue distances and their statistical significance. Applying these measures to IDP ensembles and to a protein containing both structured and intrinsically disordered domains provides deeper insights into how structural features relate to function.

## Introduction

Comparing the high-resolution structures of proteins is critical for understanding their function and evolutionary history (1, 2). Structural comparisons rely on quantitative similarity measures. The most common measure is the root-mean-square deviation (RMSD) of the atomic positions between two structures, which is minimized upon rigid-body superimposition of these structures (3, 4). But the RMSD is often not very informative as it averages out differences across regions of the structures with varying similarity levels. Therefore, superimposition-independent measures relying on inter-residue distances have been proposed, which are moreover invariant under reflection unlike the superimposition-based RMSD (5). Distance-based metrics have been used to compare well-defined protein structures (4–6), simulated or experimentally restrained conformational ensembles (7–10) or unfolded states of proteins (11–13).

Similar to the unfolded state, intrinsically disordered proteins (IDPs) and regions (IDRs) must be described as ensembles of heterogeneous, rapidly interconverting conformations. Characterizing and comparing such ensembles is therefore particularly challenging. Employing restraints primarily obtained by nuclear magnetic resonance (NMR) and small-angle X-ray scattering (SAXS), conformational ensembles have been characterized for many functionally important IDPs, such as α-synuclein (14), Sic1 (15), p27^Kip1^ (16), and tau (17), and are made accessible in the PED database (18). Owing to their high conformational variability, however, adequately characterizing an IDP/IDR ensemble from a limited amount of experimental data is an inherently underdetermined problem (19). A given disordered protein may therefore be modeled as multiple, seemingly equivalent ensembles, representing alternative fits to the experimental data (18). Although these alternative ensembles may carry functionally relevant structural information (20, 21), their critical analysis and comparative evaluation is particularly challenging and has so far not been attempted for two main reasons. First, their extreme conformational heterogeneity makes it difficult to evaluate the degree of global similarity between two ensembles by any measure, let alone by RMSD-based metrics. Second, the function of disordered proteins is often mediated by short, sequentially contiguous binding motifs (22, 23) adopting locally relevant conformations. The latter are interconnected through more structurally variable linkers (24) that determine the relative overall configuration of these important motifs. Therefore, the similarity of IDP/IDR ensembles must be evaluated at both the local and global levels in a statistically meaningful approach.

To address these issues, we developed superimposition-independent measures for evaluating the local and global similarity between two ensembles. The local similarity between specific regions of the polypeptide is evaluated from the differences between the distance distributions of individual residue pairs, and their statistical significance. The global similarity is captured by the *ens_dRMS* measure, an RMSD-like quantity representing the root-mean-square difference between the medians of the inter-residue distance distributions of the two ensembles. We show that our superimposition-free structural similarity measures are effective in describing both global and local differences between conformational ensembles of IDPs/IDRs derived using experimental restraints or by molecular simulations, and that they also conveniently quantify the structural similarity of proteins containing both structured and disordered regions.

## Material and Methods

### Datasets of protein conformational ensembles

#### Conformational ensembles of intrinsically disordered proteins (IDP) and regions (IDR)

Data on conformational ensembles of the fully disordered K18 segment of human tau protein (130 residues) and the Measles virus N-tail protein (132 residues,(17)) were downloaded from the Protein Ensemble Database (PED) (18), which currently stores such data for 16 different protein systems or fragments thereof, comprising more than 50 ensembles of 24 fully or partially disordered protein regions. For both tau-K18 and MeV N-tail, 5 ensembles comprising 199 conformations were retrieved. These ensembles were generated by combining conformational sampling with ensemble selection based on NMR data from residual dipolar coupling (RDC) and chemical shift (CS) analyses (17, 25). Random-pool ensembles for the two systems, (comprising 100 ensembles of 200 conformers each), were obtained from the authors (Martin Blackledge personal communication). These random pools were generated as previously described (25).

#### Conformational ensembles generated using molecular dynamics simulations with different force fields

Five distinct ensembles of the intrinsically disordered 24-residue serine/arginine (SR)-rich peptide (residues 22-45 of SR-rich splicing factor 1) were generated using μs time scale replica exchange molecular dynamics (MD) simulations (26) with GROMACS 4.5.4(27) using CHARMM(28) (CHARMM22*(29), CHARMM36(30)) and AMBER(31) (99sb*-ildn(29), 03w(32)) force fields, moreover, with CAMPARI using the ABSINTH implicit solvent model(33). Here, using simulation parameters identical to those previously described(31), with GROMACS 2016.3 and the CHARMM22* force field with CHARMM-modified TIP3P water, three independent high temperature (600 K) 0.2 μs long MD simulations were used to generate a model for the random coil ensemble of the same system, denoted as the “High-T" ensemble.

#### Conformational ensembles of the human prion protein

Three distinct conformational ensembles of truncated human prion protein (huPrP, full-length: 231 amino acids), determined by Nuclear Magnetic Resonance (NMR) were downloaded from the Protein Data Bank (PDB)(34). These were 2 huPrP(90-226) structures: 2lsb (35), 5l6r (36); and one huPrP(90-231): 5yj5 (37).

### Distance-based metrics for comparing conformational ensembles

To compare two conformational ensembles A and B, we use metrics based on the Cα-Cα distances within individual conformers in the ensembles. Since IDRs of proteins display very diverse conformations, Cα-Cα distances of a given pair of residues *i,j* of the polypeptide follow a distribution of values across conformers. This distribution differs between residue pairs and may therefore provide useful information on the variation of the spatial proximity of specific regions along the polypeptide. To derive this information, two matrices are computed for each of the two ensembles (**Figure 1, and Figure 2A)**. One is a matrix whose elements represent the median of the Cα-Cα distance distributions *dμ(i,j)* for equivalent residue pairs *i,j* in the conformations belonging to the same ensemble. The other matrix contains the standard deviations *dσ(i,j)* of the corresponding distributions.

**Figure 1.**
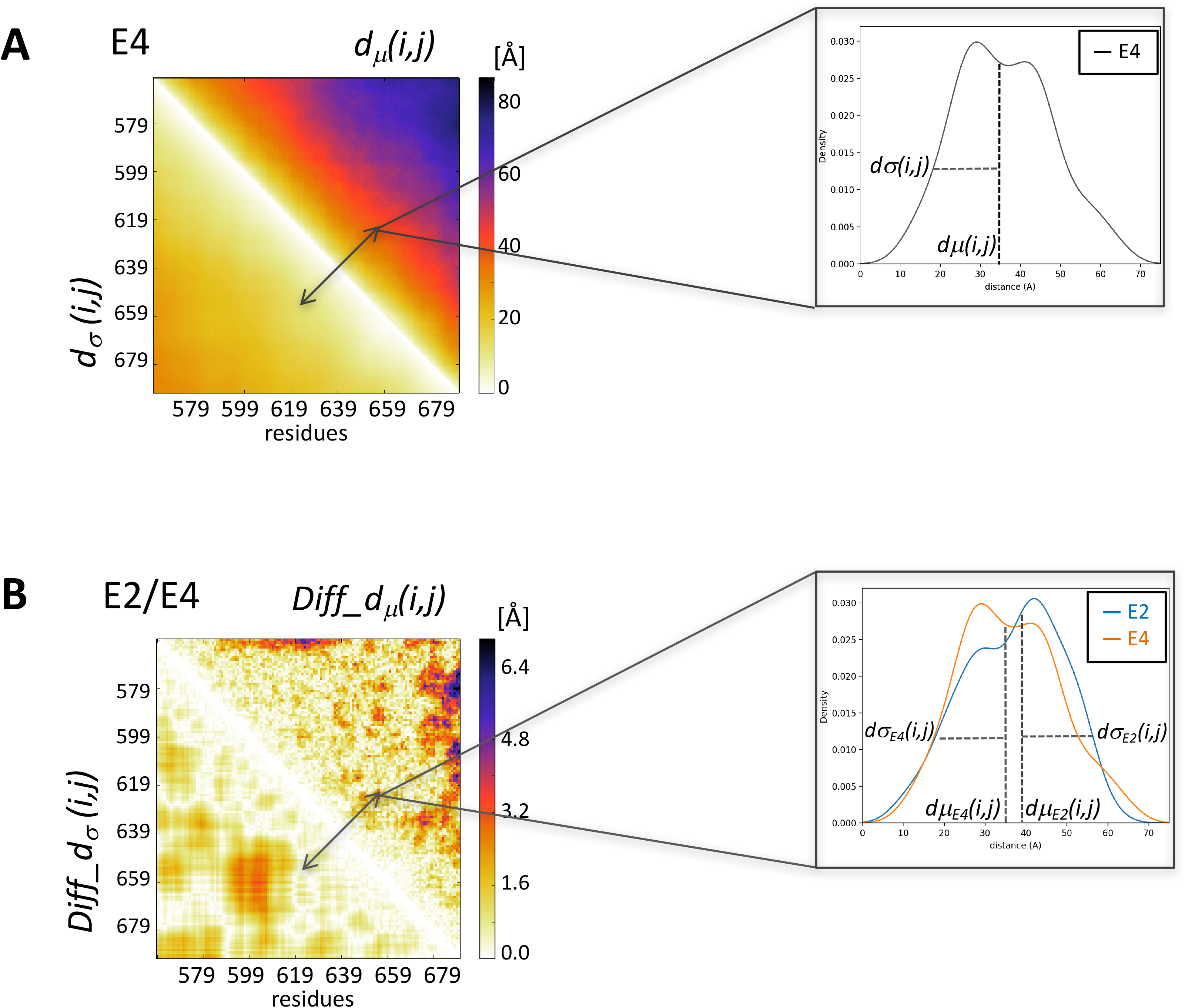
Composite heatmaps representing the *dμ(i,j)/ d*σ*(i,j), and Diff_dμ(i,j)/ Diff_dσ(i,j*) matrices. Upper left: Heatmap, where the upper triangle displays the median of the inter-residue distance distributions *dμ(i,j)* (computed between Cα atoms) for equivalent residue pairs *i,j* in the conformations of one ensemble (E4 of the MeV N-Tail domain). The lower triangle displays the standard deviations *d*σ*(i,j)* of the corresponding distributions. Upper right: example of the distribution contributing to one element of the *dμ(i,j)/ d*σ*(i,j) composite heatmap.* Lower left: Heatmap where the upper triangle depicts the *Diff_dμ(i,j)* matrix whose elements are the absolute differences of the medians of the Cα-Cα distance distributions between two ensembles (E2/E4 of the MeV N-Tail domain); the lower triangles displays the absolute differences between the corresponding standard deviations *Diff_dσ(i,j*). Lower right: example of 2 distance distributions contributing to one element of the *Diff_dμ(i,j)/ Diff_dσ(i,j*) composite heatmap.

**Figure 2.**
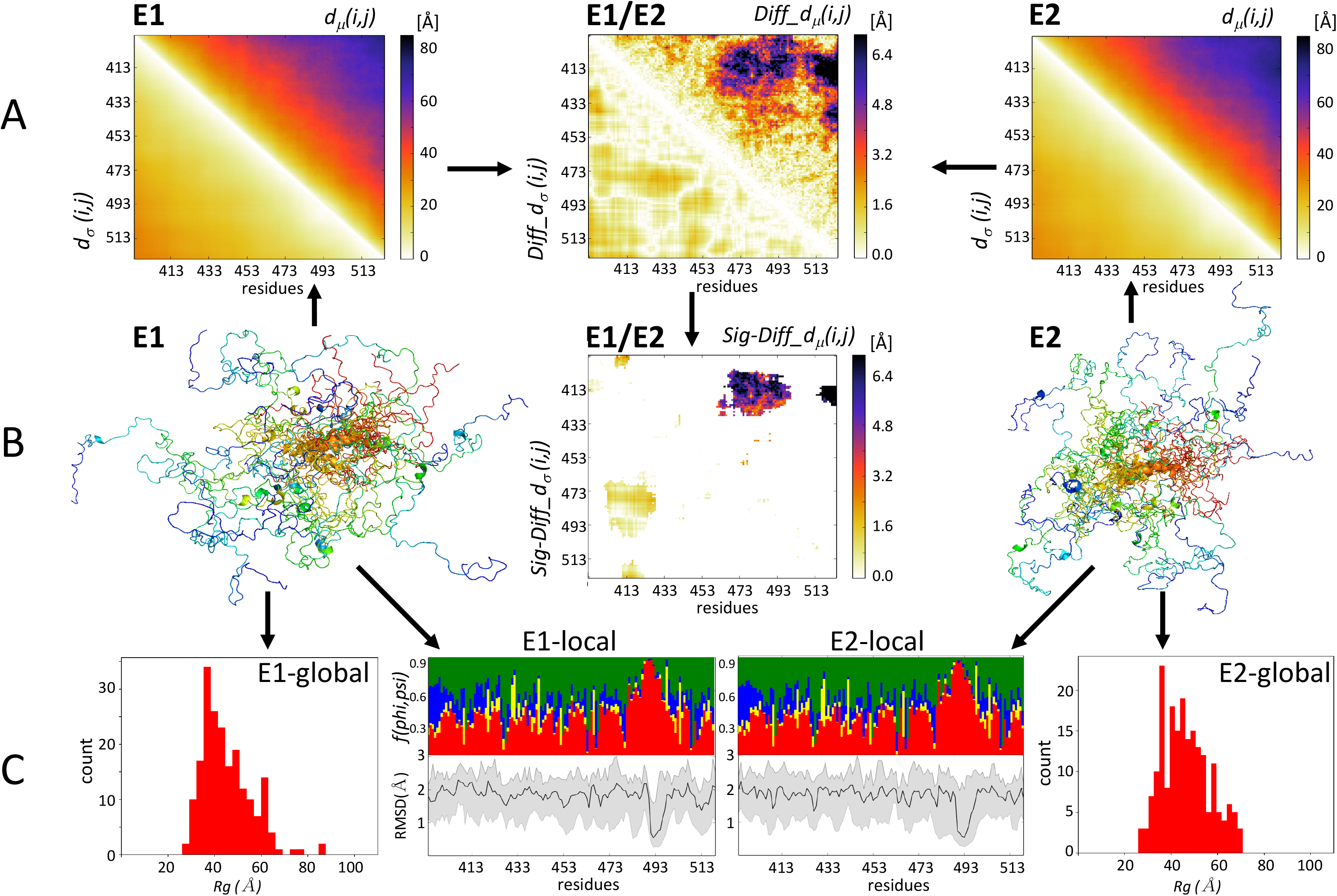
The ensemble comparison approach: an overview. (A) Displays 3 heatmaps. Two represent a composite matrix for each of the ensembles *E1* and *E2* that are being compared. The upper triangle of each map displays the medians of the inter-residue distance distributions *dμ(i,j)*. The lower triangle displays the standard deviations *d*σ*(i,j)* of the corresponding distributions. The third heatmap depicts the composite difference matrix (*E1/E2*) between the two ensembles; its upper triangle depicts the *Diff_dμ(i,j)* matrix whose elements are the absolute differences between the *dμ(i,j) distributions* in the two ensembles; its lower triangles represents the absolute differences between the corresponding standard deviations *Diff_dσ(i,j*) (see **Materials and Methods** for detail). Inter-residue distances and the differences thereof are given in Angstroms (Å). (B) Heatmap highlighting only *Sig_Diff_dμ(i,j)* for the *E1/E2* pair, i.e. elements of the *Diff_dμ(i,j)* matrix representing statistically significant differences between the corresponding *d(i,j)* distributions (p<0.05) (upper triangle), and the corresponding *Diff_dσ(i,j*) values (lower triangle). The highlighted differences concern *d(i,j)* distance distributions between a segment spanning residue 473-493 and the N-terminal region (residues 405-420) of the polypeptide. Flanking this map are cartoon models depicting the backbones of individual conformations of the *E1* (left) and *E2* (right) ensembles. The conformations are color coded from blue (N-terminus) to red (C-terminus) and are superimposed onto the equivalent helical segment (orange ribbon) in both ensembles. (C) Shown are graphs of the local flexibility properties and conformational preferences of the *E1, E2* ensembles (*E1-local; E2-local*). The lower graph plots of the medians and 95 percentile confidence interval of the local backbone RMSD distributions for conformations of the E1 and E2 ensembles. The upper graph shows the fraction of the conformations in each ensembles adopting backbone (ϕ,Ψ) values mapped onto the corresponding 4 regions of the Ramachandran map (**Supplementary Figure S10**; red: α-helix; yellow: left-handed helix; blue: β-strand; green: poly-proline I/II) (17). The RMSD values are computed as described in **Materials and Methods**. The local plots are flanked by bar graphs showing the *Rg* distributions of conformations in each ensemble (*E1-global, E2-global*), which indicate that the E2 ensemble is somewhat more compact than E1.

For each pair of conformational ensembles A and B, we then compute a difference matrix (**Figure 1 bottom, and Figure 2A, center**), where the *i,j* elements above the diagonal represent the absolute values of the difference between the median distances between residues *i,j*:

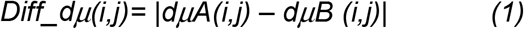

whereas the *i,j* elements below the diagonal contain the absolute differences between the corresponding standard deviations:

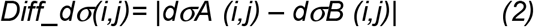

Since the difference matrix evaluates differences of *i,j* distance distributions, it is important to assess the statistical significance of these differences (**Figure 2B, center**). This is done using the non-parametric Mann–Whitney–Wilcoxon test, so that the resulting difference matrix displays *Diff_dμ(i,j),* and *Diff_dσ(i,j)* values only for statistically different *d(i,j)* distributions (p<0.05).

The difference matrices of Eq (1) deal with differences between median values of distances, which themselves may span a wide range sizes (from 3-20 Å). The same *Diff_dμ(i,j)* value of, say, 5 Å amounts to a more drastic distance variation for a median distance of 10 Å than for that of 50 Å. To account for this bias, we also compute normalized difference matrices:

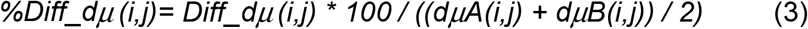

To provide a single global measure of the differences between two conformational ensembles A and B, we computed the *ens_dRMS,* defined as:

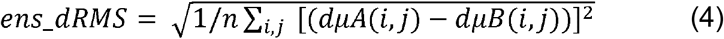

Where *dμA(i,j)* and *dμB(i,j)* are the medians of the distance distributions of *i,j* residue pairs in ensembles A and B respectively, and *n* equals the number of *i,j* pairs. The *ens_dRMS* is computed over all *i,j* pairs of the conformations in the two ensembles, to enable comparison between different ensembles of the same polypeptide.

In addition to these distance-based metrics we compute the radius of gyration, *Rg*, of the conformations in the ensemble, using only Cα atom coordinates. We use it as a global measure of the ensemble dimensions (**Figure 2C**).

Using the median of the Cα-Cα distance distributions instead of their averages as the basis for our ensemble comparison metrics has the advantage of representing a more robust measure. A significant fraction of these distances is not normally distributed, their average value may therefore be readily affected by a few outlier values. This is not the case of the median, which represents the most highly populated value. Clearly however, difference matrices computed using the two measures are closely related and our approach could readily accommodate either measure. Indeed, difference matrices obtained for pairs of experimentally derived ensembles using respectively, the median and the root-mean-square average *d(i,j)* value (or the average *d(i,j)* value) (see **Supplementary Section 1.1**), display virtually identical patterns. Nevertheless, the *Diff_dμ(i,j)* matrix exhibits more prominent features than the difference matrix based on the root-mean-square average *d(i,j)* value, in line with the more robust nature of the median, as illustrated in **Supplementary Figure S1A,B**. Global measures based on *d(i,j)* averages, computed for the 10 pairs of experimentally characterized human tau-K18 IDP/IDR ensembles, are also highly correlated (Pearson’s r = 0.87,0.89) with the *ens_dRMS* values.

We also note that measuring differences in distances between backbone atoms (here Cα-Cα distance) in IDP ensembles is a reasonable first step. In these ensembles, including those generate by molecular dynamics simulations, side chains remain highly flexible. They sample, on average, a much wider range of orientations than in globular proteins; and hence, their interactions are in general weaker. In support of this view, we find that considering Cβ-Cβ distance distributions instead of those between Cα-Cα atoms yields indistinguishable results, as illustrated in the **Supplementary Figure S2A,B.** When dealing with proteins containing structured regions, or when formation of more long-lived contacts with side chains are of interest, our backbone-centric approach would need to be refined to quantify differences in contacts between both side chains and backbone atoms.

### Relation to other metrics for comparing ensembles

#### Metrics for quantifying global differences between ensembles

The *ens_dRMS* of Eq (4) is related to, but distinct from, several other intuitive or published metrics that quantify the global difference between two conformational ensembles.

One obvious metric is the difference between the average *Rg* values of the conformations in each of the two ensembles that are being compared. Another example is the difference between inter-residue distance-based version of the ensemble “structural radius” originally defined using position-dependent RMSD (38). The latter is expressed as the root-mean-square average pairwise RMSD between all pairs of conformations in a given ensemble and captures the structural diversity of the ensemble.

Using the 5 experimentally characterized IDP/IDR ensembles of respectively, the human tau-K18 and MeV N-tail proteins of our dataset (representing 10 pairwise comparisons for each system), we evaluated the relationships between these two metrics and the *ens_dRMS* of Eq (4). The difference between average *Rg* values of two ensembles was computed as: *Diff_ensRg = |<Rg(A)> - <Rg(B)>|*, where < > indicate ensemble averages, and A, and B are different ensembles. The derivation of the distance-based version of the “structural radius” for each ensemble, *dR_struct_*, is provided in the Supplementary Material. The difference between d*R_struct_* values of two ensembles was computed as: *Diff_dR_struct_ = |dR_struct_(A) - dR_struct_(B)|.*

This analysis revealed a moderately high correlation between the *ens_dRMS* and *Diff_ensRg* values (Pearson’s r = 0.69), computed for the same pairs of ensembles, but a very low correlation of *ens_dRMS* with *Diff_dR_struct_* (**Supplementary Figure S3A,B**). The low correlation with *Diff_dR_struct_* was mainly due to the very low correlation between these two quantities for the 10 pairs of the MeV N-tail ensembles, caused by the more nature of one of the ensembles (**Supplementary Table S1, Supplementary Figure S3D**). These results confirm that the *ens_dRMS,* is indeed a distinct measure from metrics such as *Diff_ensRg* and *Diff_dR_struct_*. The *ens_dRMS* directly computes averages over the difference in median distances of individual residue pairs in conformations from different ensemble*s*. The other two metrics first compute averages over distances within conformations within the same ensemble (*Rg*), or over the difference in distances between conformations, again within the same ensemble (*dR_struct_*). Both metrics then use these ensemble averages to quantify the between-ensemble differences. Interestingly we find that *Diff_ensRg and Diff_dR_struct_,* are only moderately correlated with one another (Pearson’s r = 0.53), indicating in turn, that even these seemingly related measures quantify distinct average global features of the conformational ensembles (**Supplementary Figure S3C**).

#### Metrics based on the Kullback–Leibler divergence (KLD) of two distributions

Our distance-dependent metrics evaluate quantities that capture differences between the distance distributions of two ensembles, but do not evaluate the differences between the distributions themselves. A classical measure of the difference between two distributions is the *Kullback–Leibler divergence* (*KLD)* (39).

Several studies have illustrated the effectiveness of *KLD*-based metrics in comparing conformational ensembles of globular proteins generated by molecular simulations and modelled using experimental restraints (8, 40, 41). These studies analyzed distributions of different structural parameters (e.g. pairwise global RMSD values between conformations, or backbone and side chain dihedral angles) and pre-processed the underlying conformational ensembles in different ways. They also indicate clearly that extracting statistically significant values from such *KLD*-based metrics does require a large sample size and is computationally intensive. Given the small size of the experimentally restrained IDP datasets analyzed here, it is difficult to compare our distance-dependent local (*Diff_dμ(i,j), Diff_dσ(i,j)*) and global (*ens_dRMS*) metrics with *KLD*-based metrics, which measure differences between the underlying *d(i,j)* distributions.

We nevertheless performed a rough comparison of the symmetrized form of the *KLD* between *2* distance distributions, sym*KLD_d(i,j),* and our *Diff_dμ(i,j)* metric, for the 10 pairs of the tau-K18 IDP ensembles of our dataset (see **Supplementary Material Section 2.3** for detail). Results showed moderate correlation coefficients (Pearson’s *r =* 0.42-0.51) between the two metrics for pairs of ensembles exhibiting significantly different *d(i,j)* distributions, but a negligible correlation for ensemble pairs displaying no significantly different *d(i,j)* distributions (evaluated by the Mann–Whitney– Wilcoxon test). (**Supplementary Figure S4**). Interestingly however, a much higher correlation (Pearson’s *r* = 0.8) was obtained between the two global metrics, *ens_dRMS* and *ensKLD* values (the ensemble-averaged *symKLD_d(i,j)* values) between 2 ensembles computed for the 10 pairs of tau-K18 ensembles (**Supplementary Table S2**).

Taken together, these results suggest that *symKLD_d(i,j)* and *Diff_dμ(i,j)* capture differences between distinct aspects of individual *d(i,j)* distributions (or differ due the small sample size), and that these differences are averaged out when the global metrics are compared, hence suggesting that our simple global metric, *ens_dRMS*, captures rather well the differences between the underlying *d(i,j)* distributions of two ensembles. Clearly, such comparisons need to be repeated using a careful formulation of *KLD*-based metrics that depend on inter-residue distances, as well as larger datasets. Furthermore, such metrics may themselves be a useful addition to the toolbox of methods, enabling the analysis of a more complete range of properties of IDP ensembles than the metrics proposed here.

### Measuring local backbone flexibility and conformational biases within ensembles

To evaluate possible biases of individual ensembles towards specific local backbone conformations (α-helix, left-handed helix, extended β-strand etc.) as well as the extent of local backbone flexibility, we carry out superimpositions of backbone atoms for overlapping 5-residue segments along the polypeptide chain for pairs of conformations within a given ensemble (**Figure 2C, center**). The average and the 95 percentile confidence interval of the classical backbone RMSD values computed across all *k,l* pairs of conformations are computed for each segment, and assigned to the first residue of the segment, representing the residue number *n*, along the polypeptide serving as the segment anchor:

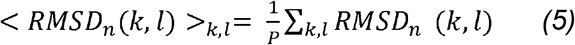

where *P* is the total number of *k,l* pairs of conformations in the ensemble.

These local backbone comparisons are complemented with an analysis of the frequencies of backbone (ϕ,Ψ) torsion angle values, mapped onto regions of the Ramachandran map corresponding to the four common secondary structure motifs: α-helix, β-strand, poly-proline, and left-handed α-helix (See (17) and **Supplementary Figure S10**).

### Code availability

https://github.com/lazartomi/ens-dRMS

## Results and Discussion

### Comparing conformational ensembles of intrinsically disordered proteins

We propose a general approach for comparing conformational ensembles that combines several complementary metrics (**Figure 2)**. At its core are novel distance-based metrics quantifying the global and local similarity between two conformational ensembles by comparing distributions of inter-residue distances.

Essential components of the distance-based metrics are two matrices for each of the ensembles E1 and E2 to be compared **(E1, E2**, heatmaps **Figure 2A**). One contains the medians of the inter-residue distance distributions *dμ(i,j)* (computed between Cα atoms) for equivalent residue pairs *i,j* in conformations of the ensemble (top right half of the heatmaps). The second contains the standard deviations *d*σ*(i,j)* of the corresponding distributions (bottom left half of the heatmaps). For a given pair of ensembles, two difference matrices are computed (**E1/E2, Figure 2A**): the *Diff_dμ(i,j)* matrix containing the absolute differences between the medians of inter-residue distance distributions of the two ensembles (top right half of the heatmaps), and the matrix containing the absolute differences between the corresponding standard deviations, *Diff_dσ(i,j*) (bottom left half of the heatmaps). Since the *Diff_dμ(i,j)* matrix evaluates differences between distributions, the statistical significance of these differences is evaluated and the resulting difference matrices list only values for the significantly different *d(i,j)* distributions (*p<0.05*) (**E1/E2 in Figure 2B**; for further details see **Materials and Methods**) named *Sig-Diff_dμ(i,j)* and *Sig-Diff_d*σ*(i,j)*.

To obtain a single global measure of the differences between the two conformational ensembles E1 and E2, we compute the root-mean-square deviation between the median distance elements *dμ(i,j)* of the conformations in the two ensembles, denoted as *ens_dRMS* (see **Materials and Methods**).

The distance-based metrics are complemented with several classical measures applied to individual ensembles (**Figure 2C,** and **Materials and Methods**). The local backbone variability within one ensemble is quantified by the distributions of the average backbone RMSD values of 5-residue segments along the polypeptide, computed over pairs of conformations in each ensemble. Local conformational preferences within a given ensemble are evaluated by the frequencies of backbone (ϕ,Ψ) torsion angles of individual residues, mapped onto regions of the Ramachandran map corresponding to secondary structure motifs (see **Materials and Methods**). We see for example, that the region near residue 490 of the polypeptide in the analyzed ensembles displays low backbone variability and adopts a helical conformation in both E1 and E2 ensembles (E1-local / E2-local, **Figure 2C**), but that the same region adopts different spatial positions relative to the N-terminus of the polypeptide in the two ensembles (residues 400-430) (E1/E2 heatmap, **Figure 2B**).

Global conformational parameters of individual ensembles are also quantified from the distribution of the radius of gyration (*Rg)* of conformations within an ensemble, with examples presented below.

### Application to conformational ensembles of specific protein systems

To illustrate the potential of our approach, we apply it to experimentally characterized IDR ensembles of two proteins. One is the N-tail region (132 residues) of the measles virus (MeV) nucleoprotein, which includes a short transient α-helix that mediates the interaction with the C-terminal X domain of the MeV phosphoprotein (42, 43), which is important for the replication of the viral genome. The second is the K18 segment (130 residues) of human tau protein, a microtubule-associated protein, which binds microtubules via 4 imperfect microtubule-binding repeats (R1-R4) located within the K18 segment (44, 45), and promotes microtubule polymerization and stability (see **Supplementary Figure S5** for the sequences of these domains). For each of these proteins, 5 ensembles comprising 199 conformations were retrieved from the PED database (see **Materials and Methods**). These ensembles represent distinct modeling solutions derived by sampling random coil conformations, denoted here as ‘random pool’, followed by ensemble selection based on the fit to NMR data (residual dipolar coupling and chemical shift data) as described in refs (17, 25).

The *ens_dRMS* values for the 10 pairs of conformational ensembles of the tau-K18 and MeV N-tail segments (**Table 1**) span a very similar small range: 1.47-2.15 Å (tau-K18) and 1.48-2.90 Å (MeV N-tail), suggesting a substantial average structural similarity between the conformations of the 5 ensembles of each protein. This similarity was also reflected by indistinguishable *Rg* distributions of the conformations in the corresponding ensembles.

**Table 1:** *ens_dRM*S value for pairs of ensembles of the tau-K18 and MeV N-tail IDR.

| tau-K18 |  |  |  |  | MeV N-tail |  |  |  |  |
| --- | --- | --- | --- | --- | --- | --- | --- | --- | --- |
| *ens_dRMS* | E2 | E3 | E4 | E5 | *ens_dRMS* | E2 | E3 | E4 | E5 |
| E1 | 1.91 Å | 1.72 Å | 1.83 Å | 1.98 Å | E1 | 2.83 Å | 2.90 Å | 2.43 Å | 1.62 Å |
| E2 |  | 2.15 Å | 1.84 Å | 1.93 Å | E2 |  | 1.74 Å | 1.48 Å | 2.10 Å |
| E3 |  |  | 1.47 Å | 2.06 Å | E3 |  |  | 1.82 Å | 2.14 Å |
| E4 |  |  |  | 2.04 Å | E4 |  |  |  | 1.75 Å |

To evaluate how conformational properties between ensembles differ, we examine pairs of ensembles in each protein system featuring the largest and smallest *ens_dRMS* values in **Table 1**. For each of these pairs, we examined the *dμ(i,j)* matrices, as well as the difference matrices *Diff_dμ(i,j) and Diff_dσ(i,j)*. Results show that the *Diff_dμ(i,j)* matrix for the least similar E2/E3 pair of tau-K18 (*ens_dRMS* =2.15 Å) features 4 regions with the largest *Diff_dμ(i,j)* values (>4.8 Å) (**Figure 3, Panel I**). Only three of these regions (residues 578-582/619-622; 585-600/630-660; 620-640/660-680) represent statistically significant differences of the distance distributions between segments at medium separation (30-40 residues) along the polypeptide. In contrast, the *Diff_dμ(i,j)* matrix of the most similar E3/E4 pair of tau-K18 (*ens_dRMS* =1.47 Å) features only very small regions with *Diff_dμ(i,j)* values >4.8 Å (**Figure 3 panel II**), none of which represent statistically significant differences between the underlying distance distributions, indicating that the E3 and E4 ensembles are indistinguishable at this level of the analysis. This was the only pair of tau-K18 ensembles with indistinguishable distance distributions. All the remaining pairs (including E2/E3) display varying patterns of significant differences, but only between residues positioned at medium to large separation (20-70 residues) along the polypeptide (**Supplementary Figure S6**).

**Figure 3.**
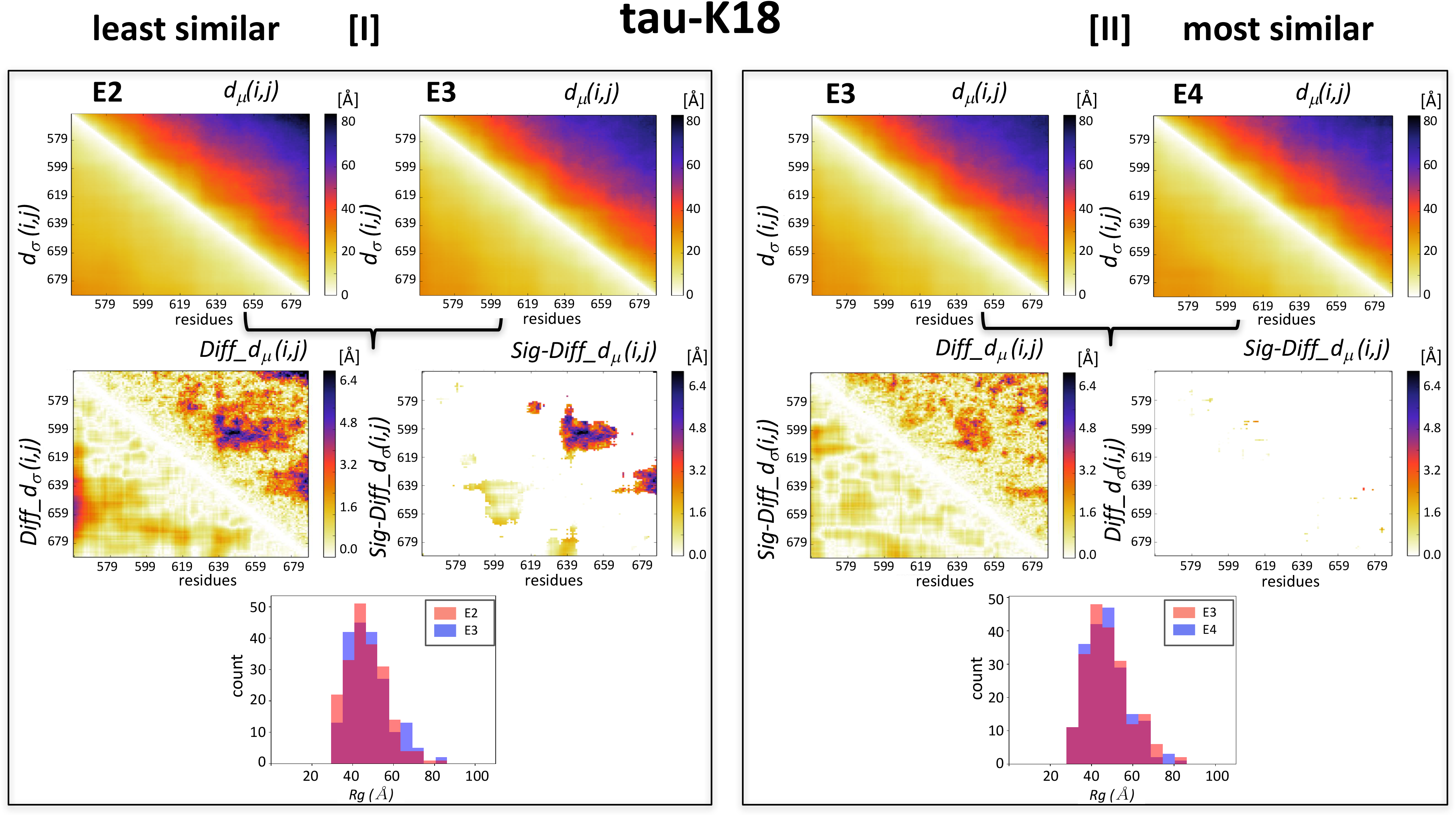
Comparisons of experimentally characterized tau-K18 IDR ensembles. Illustrated are the similarities between the E2/E3 and E3/E4 pairs of tau-K18 ensembles displaying, respectively, the largest (2.15 Å) and smallest (1.47 Å) *ens_dRMS* value in **Table 1**. Panel I: Top: heatmaps of *dμ(i,j)/d*σ*(i,j)* matrices for the individual *E2* and *E3* ensembles. Middle left: heatmaps of the *Diff_dμ(i,j)/ Diff_dσ(i,j*) computed for the E2/E3 pairs, featuring 4 regions with largest differences (>4.8 Å). Middle right: heatmaps depicting only the statistically significant elements of these maps (*Sig*-*Diff_dμ(i,j)/Sig-Diff_dσ(i,j*)). These elements span 3 regions (residues 578-582/619-622; 585-600/630-660; 620-640/660-680) representing distances between segments with medium range separation (30-40 residues) along the polypeptide. Bottom: histogram of the distributions of the gyration radii (*Rg)* of E2 and E3, found to be statistically indistinguishable (*p=0.*3). Panel II displays results for E3/E4 pair. The top, middle, and bottom panels display the same quantities as in Panel I, computed for this most similar pair. The *Diff_dμ(i,j)* and *Diff_dσ(i,j*) matrices computed for this pair highlight similar differences to those of the E2/E3 pair, but these differences are not statistically significant, resulting in the virtually empty, *Sig*-*Diff_dμ(i,j)/Sig-Diff_dσ(i,j*) heatmap. The *Rg* distributions of E3/E4 pair (bottom plot) are likewise statistically indistinguishable (*p=0.9*).

Essentially the same observations were made for the 5 ensembles of the N-tail region of MeV nucleoprotein (**Supplementary Figures S7-8**), although among the 5 N-tail ensembles, members of two pairs (E2/E4; E1/E5) were statistically indistinguishable.

These results suggest that the 5 experimentally derived conformational ensembles of the two IDP/IDR domains adopt closely similar local structures but display significant differences in their non-local structures, i.e. how short segments located at medium to large separations along the polypeptide are positioned relative to each other. Considering that the NMR data used to model the ensembles provide only local-structure restraints (17), the observed differences in non-local structure likely represent random ‘noise’ of the IDP/IDR ensemble solutions, which is not functionally relevant, and they contribute little to the average global conformational properties. This is a reasonable assumption considering that the function of IDPs/IDRs tends to be mediated by short recognition motifs that are interspersed between longer flexible linker regions adopting highly variable conformations (24), as will be further discussed below.

#### Experimentally derived versus random pool ensembles

To further characterize the experimentally derived and apparently very similar ensembles, it is important to evaluate how they differ from the ensembles of random coil conformations (random pools) from which they were selected based on the NMR data. To this end we combined all the conformations from respectively, the 5 tau-K18 and MeV N-tail ensembles (199 conformers x 5 ensembles per protein), and compared them to those of the random pools of each protein (100 ensembles x 200 conformers) generated by the authors of the ensembles (17).

Results show that the experimental ensembles of the tau-K18 and MeV N-tail proteins display similar average conformational parameters to those of their random pool versions. The two types of ensembles feature somewhat distinct *Rg* distributions for tau-K18 (*p=0.08,* **Figure 4A**), but indistinguishable distributions in the case of MeV N-tail (*p=0.4*; **Figure 4B**). The differences in the ens_dRMS distributions between the experimental and random pool ensembles (**Figure 4C,D**) are more noticeable, (although not statistically significant (p~ 0.4), due to the small sample size: only 5 experimental ensembles for each system). Those of the experimental ensembles span a narrower range (1.5-2.2 Å for tau-K18 and 1.5-2.9 Å for MeV N-tail), than their random pool counterparts (1.0-3.0 Å and 1.0-3.4 Å respectively), with a somewhat wider range for the experimental MeV N-tail than the tau-K18 ensembles. On the other hand, pairs of conformations from both types of ensembles (experimental versus random pool) follow distinct distributions of *ens_dRMS* values from those of pool-pool pairs (p=2.8e-13 and 2.5e-3 for tau-K18 and MeV N-tail respectively). These distributions display a small shift towards higher values **(Figure 4C,D**) indicating that the conformations in the experimental ensembles tend to differ more from random pool conformations than random pool conformations among each other.

**Figure 4.**
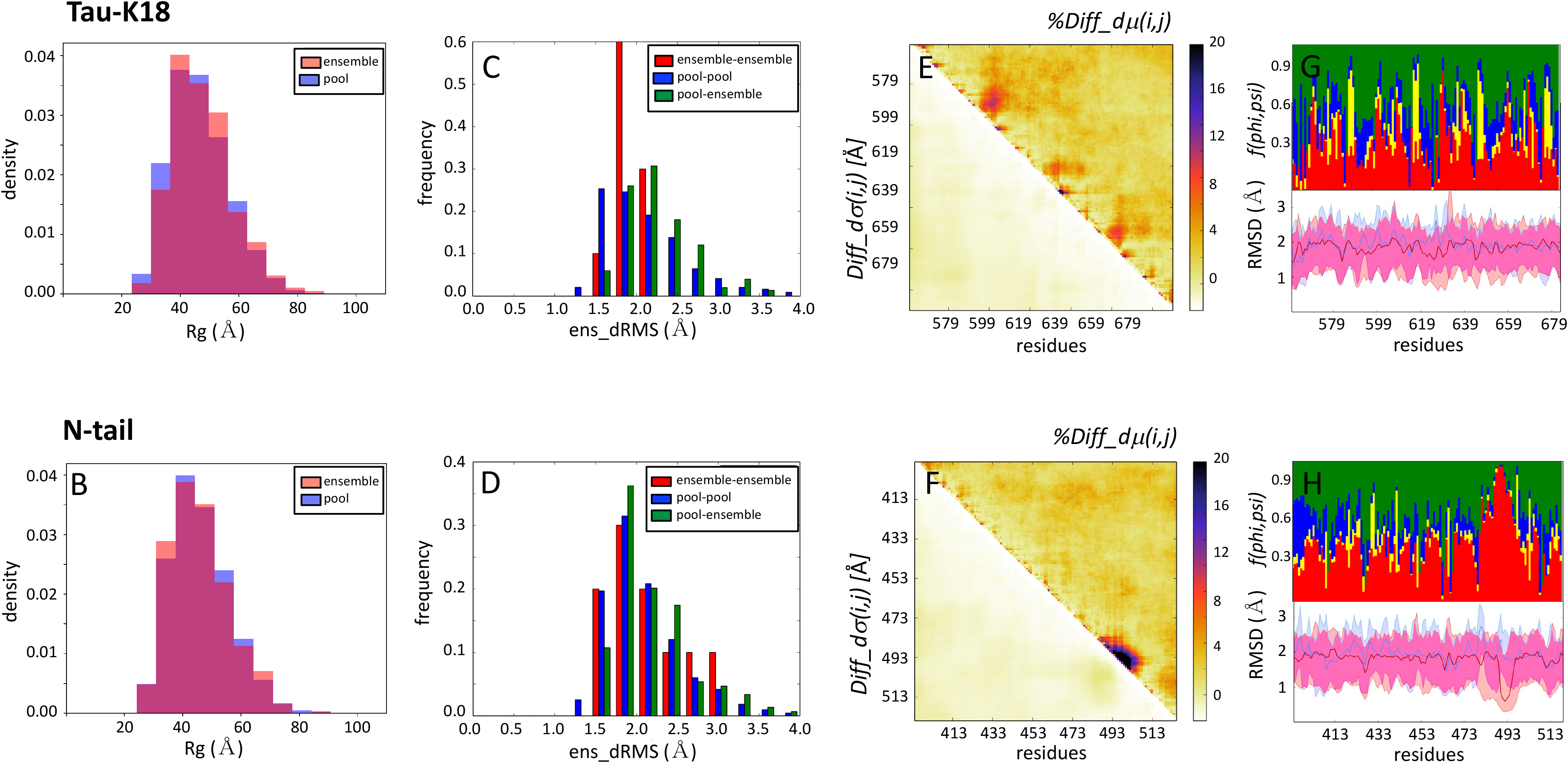
Experimental ensembles compared to their random pool versions. The conformations of respectively, all 5 tau-K18 and MeV N-tail ensembles were combined and compared to those of the random pool ensembles of each protein (25). (A), (B): Histograms of the *Rg* distributions of the experimentally restrained versus the random pool ensembles, for the tau-K18 and MeV N-tail IDR domains, respectively. The tau-K18 ensembles are somewhat more compact than their random pool version (*p=0.008*), whereas both types of ensembles are indistinguishable for MeV N-tail (*p=0.36*). (C), (D): Histograms of the pairwise *ens_dRMS* (Å) values for conformations of respectively, the experimentally restrained tau-K18 and MeV N-tail ensembles (red bars), for random pool versions of the corresponding proteins (blue bars) and for pairs comprising members of both types of ensembles (green bars). (E), (F): Heatmaps of the statistically significant portions of the normalized version of the difference matrix and the corresponding standard deviation differences, for respectively the tau-K18 and MeV N-tail IDR domains versus their random pool versions. The normalized version of the difference matrix *%Diff_dμ(i,j)*, is defined as the percent difference between the *dμ(i,j)* values from the two types of ensembles (see **Materials and Methods**). (G), (H): Twin graphs highlighting regions of the IDRs with different backbone flexibility and local structure preferences in conformations of the experimental and random pool ensembles. Bottom graph: medians and 95 percentile confidence interval of the local backbone RMSD distributions for conformations of the merged pool (blue) and experimental (pink) ensembles of respectively, the tau-K18 (G), and MeV N-tail (H) ensembles, with the overlapping portions of the plots appearing in purple. Top graph: fractions of the conformations with specific (ϕ,Ψ) preferences (see Legend of Fig. 1c for detail).

Taken together, these observations suggest that the two experimental IDP ensembles analyzed here represent conformationally biased subsets of the random pool ensemble, with the extent of bias depending on the protein system and the quality of the experimental data. It is therefore difficult to define an *ens_dRMS* threshold that reliably distinguishes experimental ensembles from their random pool versions.

To evaluate the nature of the conformational bias introduced by experimental restraints and their functional relevance, we examine differences in more local conformational parameters between the experimental and random pool ensembles.

Merging together the conformations of all 5 experimental tau-K18 ensembles (5×199 conformations), and those of the 100 random pool ensembles (100×200 conformations), we compute the distance-based difference matrices between these two sets of ensembles. **Figure 4E** plots the normalized version of the *Diff_dμ(i,j)* matrix and the differences in the corresponding standard deviations (see legend of **Figure 4** and **Materials and Methods** for detail). Interestingly, the most prominent differences are observed not between more distant regions of the tau-K18, but along the diagonal of the matrix, where 3 regions, spanning residues 580-604, 615-632, and 647-665 display significantly non-random local conformational preferences. These regions correspond to the structurally constrained microtubule-binding motifs of tau repeats (44, 45), suggesting that the differences captured by comparing experimentally restrained and random ensembles are functionally relevant. By the same logic, the significant differences in the relative positions of more distant segments often observed between the experimental ensembles are probably not functionally relevant, as already suggested above.

Local conformational preferences are likewise observed in the experimental versus random pool difference plots of the MeV N-tail domain (**Figure 4F**). They concern a contiguous segment (residues 487-507) along the diagonal, which is α-helical in the MeV N-tail ensembles but random coil in the pools. Since this helix is critical in mediating the interaction of the MeV nucleoprotein N-tail with the X-domain of phosphoprotein (42, 43), the strong signal around this motif confirms the important discriminatory power of comparing the two types of ensembles. The specific local structure preferences of the tau-K18 and MeV N-tail IDP domains are confirmed by the plots of per-residue RMSD distributions and backbone (ϕ,Ψ) values (**Figure 4G,H**).

### Comparing flexible peptide ensembles generated by molecular dynamics with different force fields

Molecular dynamics (MD) simulations are the technique of choice for modeling the dynamic properties of proteins (46), and should be a valuable tool for modeling the highly dynamic conformational states of IDPs/IDRs. For IDPs of small enough size, one may indeed expect *de novo* MD simulations to generate realistic models of conformational ensembles without experimental restraints, provided appropriate force fields are used and conformational space is sufficiently sampled (47).

This was the rationale of an earlier study of Rauscher *et al.* (26), in which μs-timescale MD simulations were run using eight different force fields and solvent model combinations to generate conformational ensembles for the intrinsically disordered 24-residue serine/arginine (SR)-rich peptide (residues 22-45 of SR-rich splicing factor 1). These ensembles were then evaluated for their consistency with NMR chemical shifts, scalar couplings, and hydrodynamic radius derived from SAXS data, measured for the same system. This comparison allowed the authors to identify the force fields that produced ensembles that were in agreement with the experimental data (26).

Here we illustrate how our ensemble comparison measures may be used to obtain useful insights into the differences between ensembles of the (SR)-rich peptide generated using 5 different force-fields, as described in reference (26). Furthermore, we evaluate how these ensembles differ from a ‘random’ ensemble generated by MD simulations of the same system at high temperature (600 K) using the CHARMM22* force field with the CHARMM-modified TIP3P water model, denoted as the “High-T” ensemble (see **Materials and Methods** for details).

Analysis of the *Rg* and *ens_dRMS* values of the different ensembles (**Table 2** and **Supplementary Figure S9**) confirms earlier findings (26) that the Amber-99sb*-ildn and CHARMM36 force fields produce the most compact ensembles. These ensembles are shown here to differ most from the High-T ensemble as witnessed by the larger corresponding *ens_dRMS* values. Ensembles produced by the Amber-03w, ABSINTH and CHARMM22* force fields feature similar *Rg* distributions to those of the High-T ensemble, and the smallest *ens_dRMS* values relative to that ensemble.

**Table 2:** *ens_dRMS* values for pairs of ensembles of the (SR)-rich peptide, derived using MD simulations with five different force fields, and additionally one at High-T.

| $ens\_dRMS$ | 99sb | Absinth | c22 | c36 | High-T |
| --- | --- | --- | --- | --- | --- |
| 03w | 6.91 Å | 1.76 Å | 1.78 Å | 5.74 Å | 1.80 Å |
| 99sb |  | 8.13 Å | 5.51 Å | 2.26 Å | 7.47 Å |
| Absinth |  |  | 2.83 Å | 6.72 Å | 1.37 Å |
| c22 |  |  |  | 4.32 Å | 2.07 Å |
| c36 |  |  |  |  | 6.09 Å |

**Figure 5** illustrates the detailed results for 2 ensembles: those produced with the CHARMM22* and CHARMM36 force fields, reported as featuring, respectively, the best fit, and a poor fit to the experimental data in the original study (26). The small normalized *Diff_dμ(i,j)* values (≤10%) between the CHARMM22* and High-T ensembles (**Figure 5C**) confirm the close structural similarity between the two ensembles, also reflected by their similar wider *Rg* distributions (**Figure 5D**).

**Figure 5.**
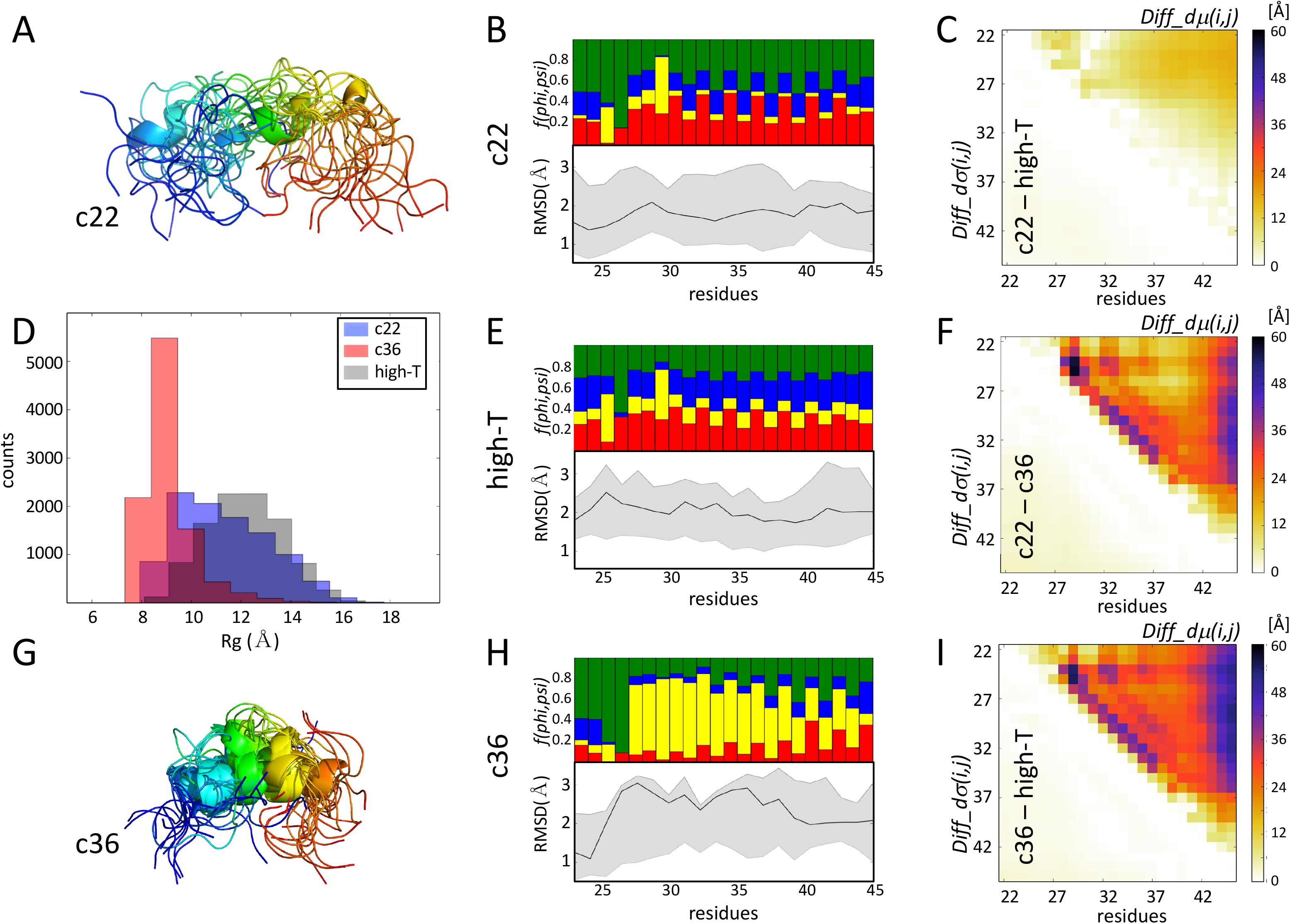
Comparing ensembles of an intrinsically disordered peptide generated using molecular simulations. Shown is a pictorial summary of the comparative analysis of 2 ensembles, generated using *de novo* molecular dynamics (MD) simulations for the 24-residue serine/arginine (SR)-rich peptide (residues 22-45 of SR-rich splicing factor 1). These ensembles were generated with the CHARMM22* and CHARMM36 force fields, previously reported as featuring respectively, the best fit, and a poor fit to the experimental data in the original study, (26). (A) Cartoon representation of the superimposed first 15 conformations from the (SR)-rich peptide ensemble generated using the CHARMM22* force field, color-coded from blue (N-terminus) to red (C-terminus). (B) Twin plots highlighting regions of the peptide with different backbone flexibility and local structure preferences in conformations of the CHARMM22* ensemble. Bottom graph: medians and 95 percentile confidence interval of the local backbone RMSD distributions for conformations of the ensemble. Top graph: fractions of the conformations with specific (ϕ,Ψ) torsion angle preferences (see Legend of Fig. 1C for detail). (C) Heatmap displaying the statistically significant portions of the *Diff_dμ(i,j)/Diff_dσ(i,j*) matrices computed between the CHARMM22* ensemble, and an ensemble generated by high temperature MD simulations (High-T), using the same force-field. This map shows only small differences, indicating a rather high degree of conformational similarity between the CHARMM22* ensemble and its High-T counterpart. (D) Histogram of the *Rg* distributions of the ensembles generated using respectively the CHARMM22*, CHARMM36 and denatured (High-T) ensembles, illustrating the higher compactness of the CHARMM36 ensemble relative to the other 2 versions. (E) Twin plots analogous to those in (B), computed for the High-T ensemble. (F) Heatmap displaying the statistically significant portions of the *Diff_dμ(i,j)/Diff_dσ(i,j*) matrices computed between the CHARMM22* and CHARMM36 ensembles. (G) Cartoon representation of the superimposed first 15 conformations from the (SR)-rich peptide ensemble generated using the CHARMM36 force field, color-coded as in (A). (H) Twin plots highlighting regions of the peptide with different backbone flexibility and local structure preferences in conformations of the CHARMM36 ensemble. (I) Heatmap displaying the statistically significant portions of the *Diff_dμ(i,j)/Diff_dσ(i,j*) matrices computed between the CHARMM36 and High-T ensembles.

Significantly larger normalized *Diff_dμ(i,j)* values, reaching up to 60%, are observed when comparing the CHARMM36 ensemble with both the CHARMM22* version and the High-T ensemble (**Figure 5F,I**). The largest differences occur between the C-terminus and residues 22-36 of the peptide, and more locally in the segment spanning residues 24-34. The latter segment is highly enriched in left-handed helix conformations in the CHARMM36 ensemble, as clearly visible on the per-residue secondary structure frequency plot (**Figure 5B,E,H).** As a result, this ensemble is also more compact (<*Rg*> = 9 Å) than the other two ensembles (**Figure 5D**). Considering the poor fit of the CHARMM36 ensemble to the experimental data, the formation of this helical structure was deemed an artifact of the CHARMM36 force field in the original study.

Thus, when ensembles are modeled *de novo*, e.g. in the absence of experimental restraints, situations may arise where statistically significant differences between an ensemble and its random counterpart have no physical or functional relevance, but merely reflect biases introduced by the modeling procedure.

### Comparing conformational ensembles of partially disordered proteins

The PDB contains many examples of proteins that feature a mix of structured domains and IDRs. These are mainly smaller proteins whose structures are determined by NMR and represented as conformational ensembles often spanning both the structured and disordered domains. Comparing such ensembles is challenging. It often involves structural superimpositions of the structured domains, which are then used to derive positional backbone and side chain fluctuations for the superimposed residues, whereas the IDRs are usually described only qualitatively.

Here we show that our ensemble-comparison protocol enables a quantitative description of such systems, which provides deeper insights into the structure-function relationship than analyses based on classical RMSD values. As an example, we use the human prion protein (huPrP), considered the causative agent of diverse prion diseases in humans, such as Creutzfeldt-Jakob disease (CJD), kuru and fatal insomnia (FI) (48). huPrP can undergo an autocatalytic, self-templated structural rearrangement to an infectious, transmissible scrapie state that causes propagation of the disease (49). The protein features an intrinsically disordered N-terminal domain (1-125) and a folded, α-helical C-terminal domain (126-226) (35). We analyze three NMR ensembles of huPrP with 20 conformers each, downloaded from the PDB. Two are of a construct of huPrP(90-226) (2lsb(35), 5l6r(36)) containing the folded domain and the adjoining 35 residues of the disordered domain, and one is for a somewhat different construct of huPrP(91-231) (5yj5 (37)).

**Figure 6** summarizes the main results for the huPrP system. It lists the average pairwise backbone RMSD values (**Figure 6**, RMSD Tables) corresponding to four different structural superimpositions of the conformations from the 3 considered ensembles. Relatively low RMSD values (1.9-2.9 Å), implying clear structural similarity, are only obtained when superimposing the C-terminal structured domain (residues 126-226, **Figure 6A**) and evaluating the corresponding backbone deviations. On the other hand, rather large RMSD values, indicative of low structural similarity, are achieved for the full huPrP fragment (90-226) after superimposing its backbone (~16 Å (**Figure 6C**), or only the backbone of the structured domain (~26 Å) (**Figure 6B**). However, somewhat lower RMSD values (9-13 Å) obtained comparing only the N-terminal IDR segment (90-125) (**Figure 6D**) reflect some structural similarity for this segment, which is completely blurred by large variations of its orientation relative to the structured domain when considering the full huPrP fragment (**Figure 6B, C**).

**Figure 6:**
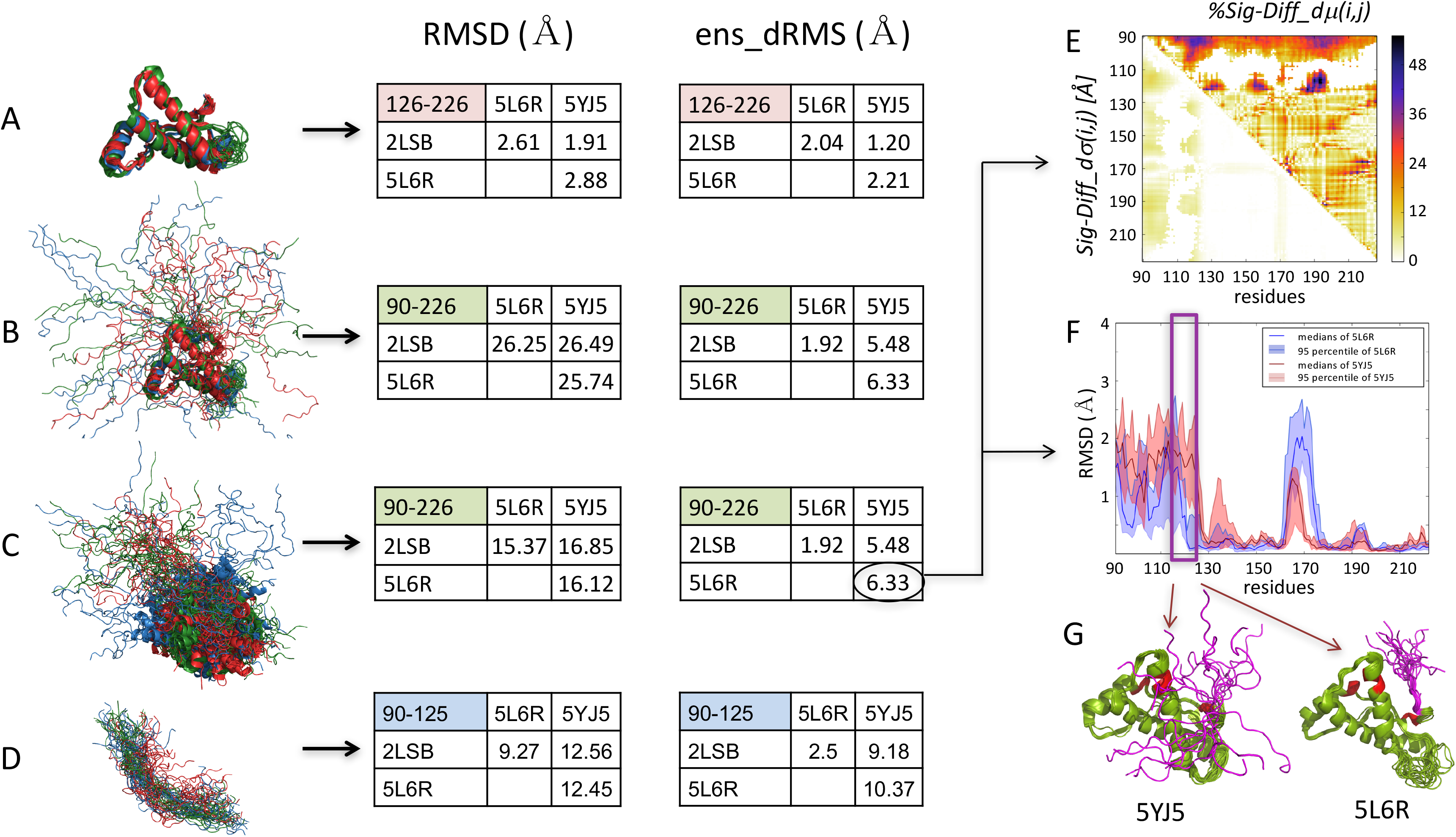
Comparing ensembles of truncated human prion protein huPrP, comprising both a structured and an intrinsically disordered region. Results are displayed for the superimposed backbone structures of the 3 NMR ensembles, with 20 conformations each. Two of huPrP (90-226), (PDB codes: 2lsb, 5l6r,) and one of the huPrP (91-231) (5yj5). Also displayed are the Tables listing the corresponding *ens_dRMS* and average classical RMSD values (Å). (A,D) superimpositions and analysis were performed considering respectively only the C-terminal structured huPrP domain (residues 126-226) (A), or only the N-terminal IDR segment (residues 90-125) (D). (B) Analysis of huPrP(90-226/91-231) after superimposing the structured domain. (C) Analysis of huPrP(90-226/91-231) after superimposing the entire polypeptide (see the main text for detail). (E) The *Diff_dμ(i,j)/Diff_dσ(i,j*) heatmaps of the two huPrP NMR ensembles displaying the largest *ens_dRMS* difference (5yj5, 5lr6), depicting prominent differences involving the disordered segments (90-125). (F) medians and 95 percentile confidence interval of the local backbone RMSD distributions for conformations of the same 2 ensembles, highlighting the local differences in backbone flexibility of the 10 residue segment immediately preceding the structured domain. (G) Cartoon models of the two huPrP ensembles highlighting the different conformations of the 10-residue IDR segment relative to the C-terminal structured domain. (F) and (G) indicate that the 115-125 segment of the IDR domain features more diverse conformations in 5yj5 than in the 5l6r structure, whereas the turn segment of the structured domain (residues 160-170) adopts more diverse conformations in 5l6r.

In contrast, structural similarities between the 3 huPrP ensembles can be clearly concluded from the superimposition-free *ens_dRMS* measure, even when comparing only the IDR segment (**Figure 6**, *ens_dRMS* Tables). The *ens_dRMS* and RMSD values are comparable for the structured domain (**Figure 6A**), confirming that the distance-based measure is effective in quantitatively describing similarity of folded proteins (5). Moreover, this measure is clearly superior to the RMSD when evaluating the similarity of the full huPrP fragment (90-226), since it features low values (**Figure 6B,C**), indicating very close structural similarity, which is not recognized by the RMSD-based analysis. Importantly, the *ens_dRMS* measure also detects a clear structural relatedness of the IDR segments, particularly in the 5l6r and 2lsb PDB entries (*ens_dRMS* ~2.5 Å), which may explain why this particular pair features a significantly smaller *ens_dRMS* (~1.9 Å) than the remaining 2 pairs (~6 Å) for the full PrP fragment.

The small *ens_dRMS* values between the 5l6r/2lsb pair likely result from a bias towards similar conformational ensembles in the corresponding NMR structures, as these structures were determined by some of the same authors (35, 36). On the other hand, the larger *ens_dRMS* values for the other 2 pairs of huPrP ensembles (~9-10 Å for the IDR (**Figure 6D)** and ~5-6 Å for the full huPrP fragment (**Figure 6B,C**)) likely reflect the different conformational properties of the huPrP 5yj5 (37), to which the two other ensembles are compared.

Indeed, for the 5yj5/5l6r pair with the largest *ens_dRMS* (6.3 Å), the ensembles of the full huPrP fragments display distinct patterns of local backbone fluctuations notably in the C-terminus of the IDR domain (residues 115-125) (**Figure 6F**). This 10-residue segment, which immediately precedes the structured domain of huPrP, stands out as displaying large differences in median distances (40-50%), relative to 3 specific regions of the structured domain, in the vicinity of residues 130, 155, and 190 (**Figure 6E**). These various features are illustrated in the molecular models of **Figure 6G**. It is noteworthy that this 10-residue huPrP segment overlaps with the palindromic sequence (AGAAAAGA) thought to be critical for the transition to the scrapie form (50), and was reported to be buried in PrP fibrils (51).

## Concluding Remarks

This study presented a novel approach for evaluating the global and local similarity of conformational ensembles of the same protein employing metrics that forego superimposition of the atomic coordinates and instead compare the distributions of inter-residue distances. These metrics are based on quantities that capture the differences between the distributions of residue-residue distances and their statistical significance. Computing these quantities is inexpensive and the results can be readily interpreted by researchers with basic knowledge in molecular modeling. We showed furthermore, that our global similarity metric, the *ens_dRMS* is distinct from other inter-residue distance-dependent global similarity metrics that evaluate the radius of gyration (*Rg*), or the so-called ‘structural radius’ (*R_struct_*) (38), reformulated here using inter-residue distances (dR_struct_). The latter metrics first average intra-molecular distances of individual conformations, or compare individual conformations within ensembles, whereas the *ens_dRMS* directly averages differences between medians of individual inter-residue distance distributions across different ensembles.

The power of our simple approach was illustrated in comparative analyses of multiple ensembles of 3 different systems: those of the MeV N-tail and tau-K18 IDR segments modeled using restraints derived from NMR experiments, ensembles of the intrinsically disordered serine/arginine (SR)-rich peptide generated by MD simulations, and NMR conformational ensembles of the human prion protein (huPrP), a protein containing a structured and an intrinsically disordered domain.

Comparison of the inter-residue distance distributions within the experimentally derived MeV N-tail and tau-K18 IDR ensembles, and between these ensembles and the random pool versions from which they were selected, readily identified previously reported regions with enhanced preferences for specific local conformational features. These regions adopted a similar pattern of inter-residue distance distributions in all the experimentally derived ensembles of both systems. However, this pattern differed significantly from that adopted by the same regions of the polypeptide in the corresponding random pool ensembles. It was particularly satisfying to verify that these very regions are functionally relevant. For tau-K18 they comprise the microtubule-binding motifs, whereas for the MeV N-tail it corresponds to the helical region mediating the interaction with the X domain of MeV phosphoprotein.

This notwithstanding, the experimental ensembles were not necessarily less conformationally diverse than their random pool versions in terms of distance distributions between more remote regions of the polypeptide. The two types of ensembles also displayed similar global compactness as measured by the corresponding gyration radii (*Rg*) distributions, suggesting in turn that non-local conformational features of the experimental ensembles that are not subjected to the restraints provided by the NMR data, conserve a ‘noisy’ random-pool like character. This could potentially be remedied by the inclusion of SAXS data in the ensemble calculation (52, 53).

Our superimposition-free structural similarity measures were likewise effective in detecting the conformational biases in ensembles of the intrinsically disordered 24-residue SR-rich peptide generated by room-temperature MD simulations using different force fields and water models. One of these ensembles was previously reported as featuring a left-handed helix conformation that was incompatible with the experimental data available for this system (26). Our approach singled out this particular ensemble as the most globally compact and locally structurally constrained that differed significantly from the ensembles derived using other force fields, or from the ensemble generated by a high-temperature MD simulation (**Figure 5**).

Lastly, applying our ensemble-comparison protocol to 3 NMR ensembles of the human prion protein (huPrP) comprising both structured and intrinsically disordered regions, provided a highly informative description of this system. In stark contrast to the RMSD-based comparisons, the superimposition-free *ens_dRMS* metric revealed sizable structural similarities between the NMR ensembles of the huPrP(90-226) fragment that includes the C-terminal segment (about 35 amino acids) of the disordered PrP domain. The *ens_dRMS* also detected a particularly close structural relatedness between both the full length (90-226) and the IDR portions (90-125) in two of the huPrP structures, which, as we subsequently verified, were determined by the some of the same authors. Quite remarkably still, in the third huPrP structure, our analysis discovered significant differences in the median distances between the IDR segment around residues 115-125 and residues of the adjoining structured domain, which we could attribute to the substantial differences of the local backbone flexibility and orientation of the 10-residue disordered segment in the huPrP structures (**Figure 6E,F,G**). It was gratifying to find that this very segment is believed to be critical for the transition of huPrP to the disease-associated, scrapie form.

Our distance-based structural similarity measures should be very useful for evaluating the global and local similarity between conformational ensembles of the same or closely related IDPs/IDRs, or proteins with both structured and disordered regions. By quantifying the structural relatedness of these flexible systems, a task that classical RMSD-based analyses strain to accomplish, a deeper insight is provided into how their structural features relate to function.

## Author contributions

SW, PT, and TL designed the study; WV contributed to the study design; TL carried out the analysis; SW, PT and MG, supervised the analysis; SR contributed computed serine/arginine-rich splicing factor 1 conformational ensembles. SW, PT & TL wrote the manuscript.

## Acknowledgements

This work was supported by the Odysseus grant G.0029.12 from Research Foundation Flanders (FWO), grants K124670 and K131702 from the National Research, Development and Innovation Office (NKFIH) of Hungary, and the Spearhead grant SRP51 from VUB, Brussels, Belgium. We also acknowledge the supercomputer resources provided by Compute Canada to S.R. for the molecular dynamics simulations. S.R. is supported by an NSERC Discovery Grant.

