## Supplementary Material for "Distance-based metrics for comparing conformational ensembles of intrinsically disordered proteins"

Tamas Lazar, Mainak Guharoy, Wim Vranken, Sarah Rauscher,

Shoshana J. Wodak* and Peter Tompa*

*corresponding authors

#### Supplementary methods

##### 1. Comparison with variants of the proposed distance metrics

*1.1 Evaluating differences between the root mean square averages rather than the medians of the inter-residue distance distributions.*

We computed the difference between the root mean square average of the *d(i,j)* distributions of two ensembles, $Diff\_d_{avg}RMS(i,j)$, as follows:

$$Diff\_d_{avg}RMS(i,j) = \left|\sqrt{\frac{1}{N}\sum_{i,j} d_A\,(i,j)^2} - \sqrt{\frac{1}{N}\sum_{i,j} d_B\,(i,j)^2}\right| \qquad (1)$$

where *d(i,j)* are the distances between residue pairs *i,j, N* is the total number of conformations in the ensembles, and A and B are the two ensembles that are being compared. The above equation is analogous to Eq (1) of the main text.

This yields the following global measure for the difference between two ensembles, equivalent to the *ens_dRMS* measure of Eq (4) of the main text:

$$ens\_dRMS' = \sqrt{1/n \sum_{i,j} Diff\_d_{avg}RMS(i,j)^2} \qquad (2)$$

with *n* representing the number of *i,j* residue pairs.

Heatmaps obtained for pairs of experimentally derived IDP/IDR ensembles, using respectively, $Diff\_d_{avg}RMS(i,j)$ and $Diff\_d_{\mu}(i,j)$, of Eq (1) of the main text, displayed virtually identical patterns, as illustrated for the *E1* and *E2* ensembles of tau-K18 (**Supplementary Figure S1A,B**). For these ensembles the correlation between $d_{avg}(i,j)$ and $d_{\mu}(i,j)$ values was very high (Pearson's $r > 0.99$). The main difference was that the heatmap features computed using $Diff\_d_{\mu}(i,j)$ showed somewhat better contrast than those computed with $Diff\_d_{avg}RMS(i,j)$. This may be explained by the fact that the root mean square average values tend to be affected by a few outlier values, whereas the median values are not. The latter are more robust since they represent the most populated d(i,j) value.

A high correlation (Pearson's r = 0.89 and 0.87) was also obtained between the global measures e.g. those of *ens_dRMS* vs *ens_dRMS',* computed for the 10 pairs between 5 experimentally characterized human tau-K18 ensembles.

Virtually the same results were obtained when we simply computed the difference between the average values of the *d(i,j)* distributions of two ensembles instead of Eq (1) above :

$$Diff\_d_{avg}\,(i,j) = \left|\frac{1}{N}\sum_{i,j} d_A\,(i,j) - \frac{1}{N}\sum_{i,j} d_B\,(i,j)\right| \qquad (3)$$

where A and B are the two ensembles and *N* is the number of conformations in the ensembles.

This analysis confirms that our approach could readily accommodate measures based on the average values of the *d(i,j)* distributions, with negligible effects on the results.

*1.2 Analyzing differences between Cβ-Cβ distance distributions instead of those between Cα-Cα atoms.*

See main text for details and results illustrated in **Supplementary Figure S2A,B.**

#### 2. Comparisons to other distance dependent metrics

*2.1 Comparing differences of ensemble averaged Rg with ens_dRMS values for IDP ensembles of our dataset.*

The radius of gyration, *Rg,* of individual conformers within ensembles is computed as described in the Methods section (main text). The difference between average *Rg* values of two ensembles is computed as:

$$Diff\_ensRg = |\langle Rg(A)\rangle - \langle Rg(B)\rangle| \qquad (4)$$

where < > indicates averages over conformations in an ensemble, and A and B are different ensembles.

Using the 5 experimentally characterized IDP/IDR ensembles of respectively, the tau-K18 and MeV N-tail proteins of our dataset, representing 10 pairwise comparisons for each system, we computed the *Diff_ensRg and ens_dRMS* quantities for all 10 ensemble pairs, with results listed in **Supplementary Table S1**. Scatter plots of the *Diff_ensRg* versus *ens_dRMS* values for these ensemble pairs are shown in **Supplementary Figure S3A.**

*2.2 Comparing differences of distance dependent $R_{struct}$ values to ens_dRMS for IDP ensembles of our dataset.*

Following Kuzmanic et al. [1] the distance-dependent pairwise RMS value between two conformations/structures K and L was computed as follows

$$dRMS\,(K,L) = \sqrt{\frac{1}{N}\sum_{i,j}\;(d_K(i,j) - d_L(i,j))^2} \qquad (5)$$

where $d_K(i,j)$ and $d_L(i,j)$ are inter-residue distances of equivalent residues pairs in conformations *K* and *L*, and *N* is the total number of distances.

The distance-dependent ensemble *dRMS* is computed as follows:

$$\sqrt{< dRMS^{\,2} >} = \sqrt{\frac{1}{M}\sum_{K,L} dRMS(K,L)^2} \qquad (6)$$

where *K* and *L* are pairs of conformations and *M* is the number of such pairs.

The distance-dependent structural radius of the ensemble is computed as:

$$dR_{struct} = \frac{1}{\sqrt{2}}\sqrt{< dRMS^2 >} \quad (7)$$

and the quantity *Diff_dR*$_{struct}$ is computed as the difference between the *dR*$_{struct}$ values of the two ensembles, A and B that are being compared:

$$Diff\_dR_{struct} = |dR_{struct}(A) - dR_{struct}(B)| \quad (8)$$

Using the same dataset as in Section 1.2, we computed the *Diff_dR*$_{struct}$ *and ens_dRMS* quantities for all 10 ensemble pairs, with results listed in **Supplementary Table S2A,B**. The scatter plots of *Diff_dR*$_{struct}$ versus *ens_dRMS* for these ensemble pairs are shown in **Supplementary Figure S3B.** The scatter plots of *dR*$_{struct}$ versus *Diff_ensRg* values for individual ensembles are depicted in **Supplementary Figure S3C.**

**Supplementary Table S1:** Comparison of *Diff_ensRg, Diff_dR*$_{struct}$ and *ens_dRMS* values computed for pairs of experimentally characterized IDP ensembles of respectively, the tau-K18 and MeV N-tail protein segments.

| **Ensembles** | **Diff_*ensRg*** | ***Diff_dR*$_{struct}$** | ***ens_dRMS*** |
|---|---|---|---|
| tau_E1-E2 | 0.93 | 0.94 | 1.91 |
| tau_E1-E3 | 0.05 | 0.08 | 1.72 |
| tau_E1-E4 | 0.1 | 0.2 | 1.83 |
| tau_E1-E5 | 1.33 | 0.73 | 1.98 |
| tau_E2-E3 | 0.98 | 0.86 | 2.15 |
| tau_E2-E4 | 0.83 | 0.74 | 1.84 |
| tau_E2-E5 | 0.4 | 0.21 | 1.93 |
| tau_E3-E4 | 0.15 | 0.12 | 1.47 |
| tau_E3-E5 | 1.38 | 0.65 | 2.06 |
| tau_E4-E5 | 1.23 | 0.53 | 2.04 |
| N-tail_E1-E2 | 1.23 | 0.13 | 2.83 |

| | | | |
|---|---|---|---|
| N-tail_E1-E3 | 1.16 | 0.5 | 2.9 |
| N-tail_E1-E4 | 1.02 | 0.44 | 2.43 |
| N-tail_E1-E5 | 0.31 | 0.82 | 1.62 |
| N-tail_E2-E3 | 0.07 | 0.37 | 1.74 |
| N-tail_E2-E4 | 0.21 | 0.31 | 1.48 |
| N-tail_E2-E5 | 0.92 | 0.69 | 2.1 |
| N-tail_E3-E4 | 0.14 | 0.06 | 1.82 |
| N-tail_E3-E5 | 0.85 | 0.32 | 2.14 |
| N-tail_E4-E5 | 0.71 | 0.38 | 1.75 |

The Pearson correlations between the 20 values of the 3 different measures in **Supplementary Table S1** are: *Diff_ensRg/ens_dRMS (r=0.68); Diff_ensRg/Diff_dR$_{struct}$ (r=0.53); Diff_dR$_{struct}$/ens_dRMS (r=0.07).* The low correlation for the latter two values is due to the poor correlation for values computed for the N-tail ensembles (r=-0.14). A significantly higher correlation is obtained for the tau-K18 ensembles (r=0.66). The poor correlation for the N-tail ensembles stems from the outlier behaviour of the E1 N-tail ensemble, which features the lowest <*Rg*> value, but near average *dR$_{struct}$* value (**Supplementary Figure S3D**). By removing the 4 data points corresponding to N-tail E1, the correlation between *Diff_dR$_{struct}$* and *ens_dRMS* increases to r=0.56.

#### *2.3 Comparison with metrics based on the Kullback–Leibler divergence (KLD) of two distributions*

To compute the Kullback–Leibler divergence (KLD) of the *d(i,j)* distributions in our dataset of experimentally derived IDP ensembles we used the KLD formulation for normal distributions [2]. This is an approximation, given that only ~65% of the *d(i,j)* values are normally distributed. To quantify the difference between *d(i,j)* distributions in ensembles *A* and B, we computed the symmetrized form of the *KLD* distance distributions, *KLD_d(i,j)* as follows:

$$symKLD\_d(i,j) = (KLD\_d(i,j)(A \,||\, B) + KLD\_d(i,j)(B \,||\, A) ) / 2 \quad (9)$$

The root-mean-square *symKLD_d(i,j)* differences between 2 ensembles, the *ensKLD,* was computed as follows:

$$ensKLD = 1/n \sum_{i,j} symKLD\_d(i,j) \quad (10)$$

where *i,j* are individual residue pairs, and *n* is the number of such pairs.

Results obtained using these formulations, and applying no corrections for small sample size (*d(i,j)* distance distributions for the experimentally determined IDP ensembles of our dataset comprise only ~200 data points, representing the number of conformations in individual experimentally restrained ensembles), are illustrated in **Supplementary Figure S4 and Supplementary Table S2**.
Moderate Pearson correlation coefficients ($r$ = 0.42, 0.51) were observed between the *Diff_d*$\mu$*(i,j)* and the symmetrized *KLD* values (Eq (9)) for individual *d(i,j)* distributions of the tau-K18 ensemble pairs (such as E1/E2, and E2/E3) exhibiting significantly different *d(i,j)* distributions, as evaluated by the Mann–Whitney–Wilcoxon test **(See main text and Supplementary Figure S4A,B**). But a negligible correlation was observed between the two values for the ensemble pair E3/E4 with no significantly different *d(i,j)* distributions (see **Supplementary Figure S4C**). A rather high correlation (Pearson's $r$ = 0.81) was obtained between the *ens_dRMS*, and *ensKLD* (the ensemble average*d symKLD_d(i,j)* values of (Eq (10)) across all 10 pairs of tau-K18 ensembles (**Supplementary Table S2**).

**Supplementary Table S2:** Comparison of *ensKLD* and *ens_dRMS* values computed for pairs of experimentally characterized ensembles of the tau-K18 disordered protein segment.

| **Ensembles** | ***ensKLD*** | ***ens_dRMS*** |
|---|---|---|
| Tau_E1-E2 | 0.0089 | 1.91 |
| Tau_E1-E3 | 0.0063 | 1.72 |

| | | |
|---|---|---|
| Tau_E1-E4 | 0.0060 | 1.83 |
| Tau_E1-E5 | 0.0079 | 1.98 |
| Tau_E2-E3 | 0.0099 | 2.15 |
| Tau_E2-E4 | 0.0077 | 1.84 |
| Tau_E2-E5 | 0.0054 | 1.93 |
| Tau_E3-E4 | 0.0042 | 1.47 |
| Tau_E3-E5 | 0.0081 | 2.06 |
| Tau_E4-E5 | 0.0078 | 2.04 |

#### Supplementary figure captions

**Figure S1:**

Comparing *Diff_*$d_{avg}$*RMS(i,j)* to *Diff_*$d\mu$*(i,j)* matrices.

(A) Heat plots depicting difference matrices for the tau-K18 E1 and E2 ensembles computed using *Diff_*$d\mu$ *(i,j),* based on the medians of the *d(i,j)* distributions (upper triangle), and using *Diff_*$d_{avg}$*RMS(i,j),* based on the root-mean-square average of the *d(i,j)* distributions (lower triangle)

(B) Heat maps highlighting only the statistically significant portions of the two matrices (see Methods section of the main text for details).

**Figure S2:**

*Diff_*$d\mu$*(i,j)* matrices computed using Cα-Cα and Cβ-Cβ distance distributions.

(A) Heatmaps depicting difference matrices computed using Cα-Cα distance distributions. Upper triangle: *%Diff_*$d\mu$*(i,j)* values, representing normalized differences of the *d(i,j)* distribution means; lower triangle: *Diff_*$d\sigma$*(i,j)* values, representing differences in standard deviations of the corresponding distributions (see Methods section of the main text for details)

(B) The same plots as in (A), but with *%Diff_*$d\mu$*(i,j)* and *Diff_*$d\sigma$*(i,j)* values computed using Cβ-Cβ distance distributions.

**Figure S3:**

Scatter plots illustrating the correlations between the *Diff_ensRg*, $Diff\_dR_{struct}$ and *ens_dRMS,* quantities.

(A) Scatter plot of *Diff_ensRg* versus *ens_dRMS* values computed for the 10 pairs of the N-Tail, and tau-K18 ensembles.

(B) Scatter plot of $Diff\_dR_{struct}$ versus *ens_dRMS* values computed for the same ensemble pairs as in (A).

(C) Scatter plot of *Diff_ensRg versus* $Diff\_dR_{struct}$ values computed for the same ensembles as in (A) and (B).

(D) Scatter plot of ensemble-averaged *Rg* values, $< Rg >$, versus $dR_{strut}$ values computed for the 5 tau-K18 ensembles. The E1 outlier ensemble is highlighted with a red circle.

The Pearson correlation coefficient computed between pairs of values in each plot are listed below the corresponding plot.

**Figure S4:**

Comparison of *symKLD_d(i,j)* versus $Diff\_d\mu(i,j)$ *matrices*

Heatmaps illustrating examples of difference matrices computed for 3 pairs of tau-K18 of ensembles. Shown are matrices for the E1/E2 (A) and E2/E3 (B) pairs, with statistically significant differences *d(i,j)* distributions, and for the E3/E4 pair (C), where most of the *d(i,j)* distributions are not significantly different. The upper triangle of the heatmaps/matrices display $Diff\_d\mu(i,j)$ values (Å), and the lower triangles depict values of $KLD'\_d(i,j) = (symKLD\_d(i,j) \times 100)_2$, The latter quantity was used to increase contrast, allowing the two matrices to be depicted simultaneously using a common color scale.

The Pearson correlation between *symKLD_d(i,j)* and $Diff\_d\mu(i,j)$ values for each pair of ensembles is listed at the bottom of the corresponding heatmap.

**Figure S5:**

Amino acid sequences of the measles virus (MeV) N-tail and tau-K18 segments, whose conformational ensembles (E1-E5) were analyzed in this study.

**Figure S6:**

Heat maps highlighting the *Sig_Diff_*$d\mu(i,j)$ values for the 10 pairwise combinations of the 5 human tau-K18 ensembles (E1-E5). These values represent elements of the *Diff_*$d\mu(i,j)$ matrices corresponding to statistically significant differences between the corresponding $d(i,j)$ distributions ($p<0.05$) (upper triangle), and the corresponding *Diff_*$d\sigma(i,j)$ values (lower triangle).

**Figure S7:**

Comparisons of experimentally characterized MeV N-tail IDR ensembles.
Quantifying the similarity between the E2/E4 and E1/E3 pairs of MeV N-tail ensembles displaying, respectively, the smallest (1.48 Å) and largest (2.90 Å) *ens_dRMS* value in **Table 1**. These ensembles were generated as described in references [3,4] of the main text, by creating a very large number of random coil conformations, followed by selection of a subset of conformations (here 199 conformations) that optimized the fit to NMR data.

**Panel I:** Results for the E2/E4 pair. Top: heat maps of $d\mu(i,j)/d\sigma(i,j)$ matrices for the individual *E2* and *E4* ensembles. Middle left: heat maps of the *Diff_*$d\mu(i,j)$*/Diff_*$d\sigma(i,j)$ computed for the *E2/E4* pairs, featuring several small regions with differences >4.85Å; middle right: heat maps depicting only the statistically significant elements of these maps (*Sig-Diff_*$d\mu(i,j)$*/Sig_Diff_*$d\sigma(i,j)$), and showing none of the *Diff_*$d\mu(i,j$ elements to be statistically significant. Bottom: histogram of the distributions of the gyration radii (*Rg*) of E2 and E4, found to be statistically indistinguishable ($p=0.3$).

**Panel II** Results for E1/E3 pair. The top, middle, and bottom panels display the same quantities as in Panel I, computed for this most different pair. The *Diff_d*$\mu$*(i,j)* and *Diff_d*$\sigma$*(i,j)* matrices computed for this pair feature much more prominent difference than those of the E2/E4 pair. The statistically significant elements (*Sig-Diff_d*$\mu$*(i,j)/Sig_Diff_d*$\sigma$*(i,j)*) highlight significant differences in the distance distributions between a short N-terminal segment and a longer C-terminal region. The *Rg* distributions of E1/E3 pair (bottom plot) are likewise statistically indistinguishable (*p=0.291*).

**Figure S8:**

Heat maps highlighting the *Sig_Diff_d*$\mu$*(i,j)* values for the 10 pairwise combinations of the 5 MeV N-tail IDR N-tail ensembles (E1-E5). These values represent elements of the *Diff_d*$\mu$*(i,j)*matrices corresponding to statistically significant differences between the corresponding *d(i,j)* distributions ($p<0.05$) (upper triangle), and the corresponding *Diff_d*$\sigma$*(i,j)* values (lower triangle).

**Figure S9:**

Distributions of the radius of gyration for the ensembles of the intrinsically disordered (SR)-rich peptide generated using MD simulations with 5 different force fields (see Methods for detail).

(A) AMBER (03w)

(B) C22: CHARMM22*

(C) Absinth: CAMPARI using the ABSINTH implicit solvent model

(D) AMBER (99sb*-ildn)

(E) C36: CHARMM36

(F) High-T (high temperature)

**Figure S10:**

Secondary structure classification based on the subdivision of the Ramachandran map adapted from Ozenne et al. 2012, J. Am. Chem. Soc. [DOI: 10.1021/ja306905s].

$\alpha$: $\alpha$-helix; $\beta$: $\beta$-strand; PPI: poly-proline I/II; LH: left-handed helix.

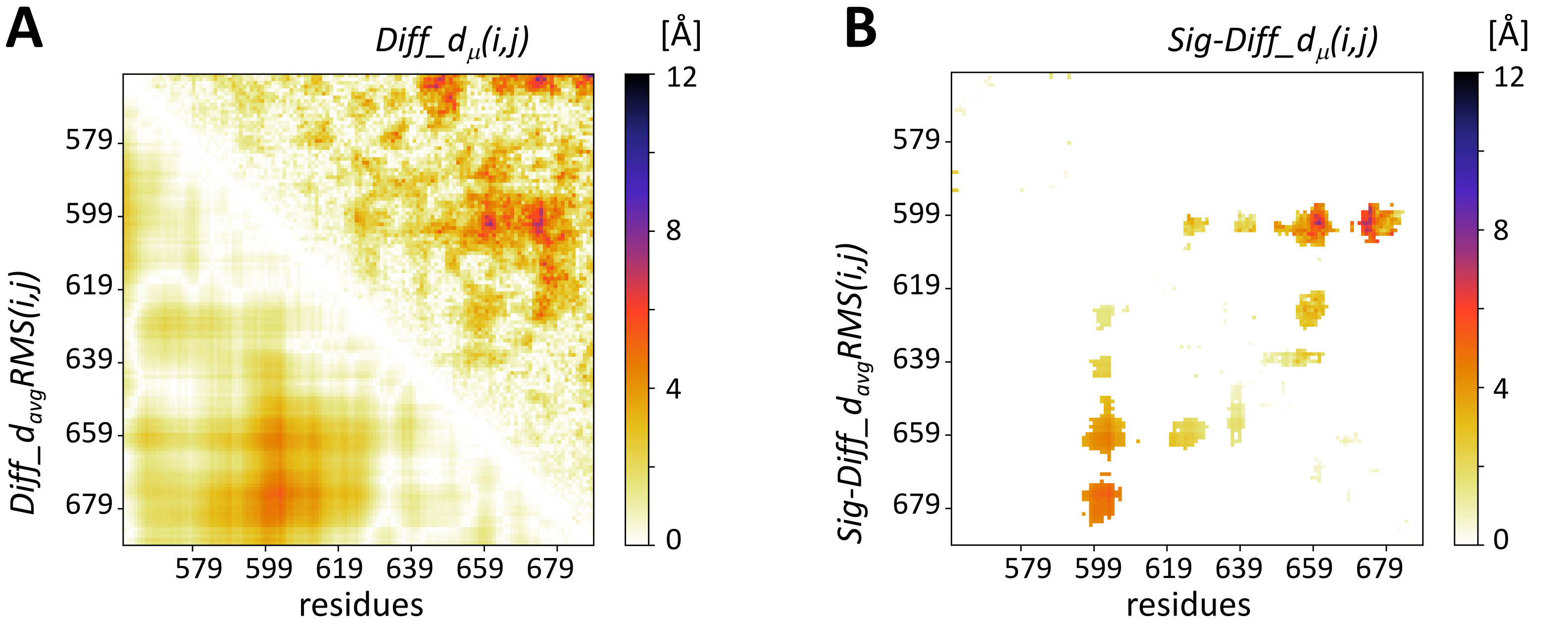

A
$Diff\_d_{\mu}(i,j)$
[Å]
12
8
4
0
579
599
619
639
659
679
$Diff\_d_{avg}RMS(i,j)$
residues
B
$Sig\text{-}Diff\_d_{\mu}(i,j)$
[Å]
$Sig\text{-}Diff\_d_{avg}RMS(i,j)$


Figure S1

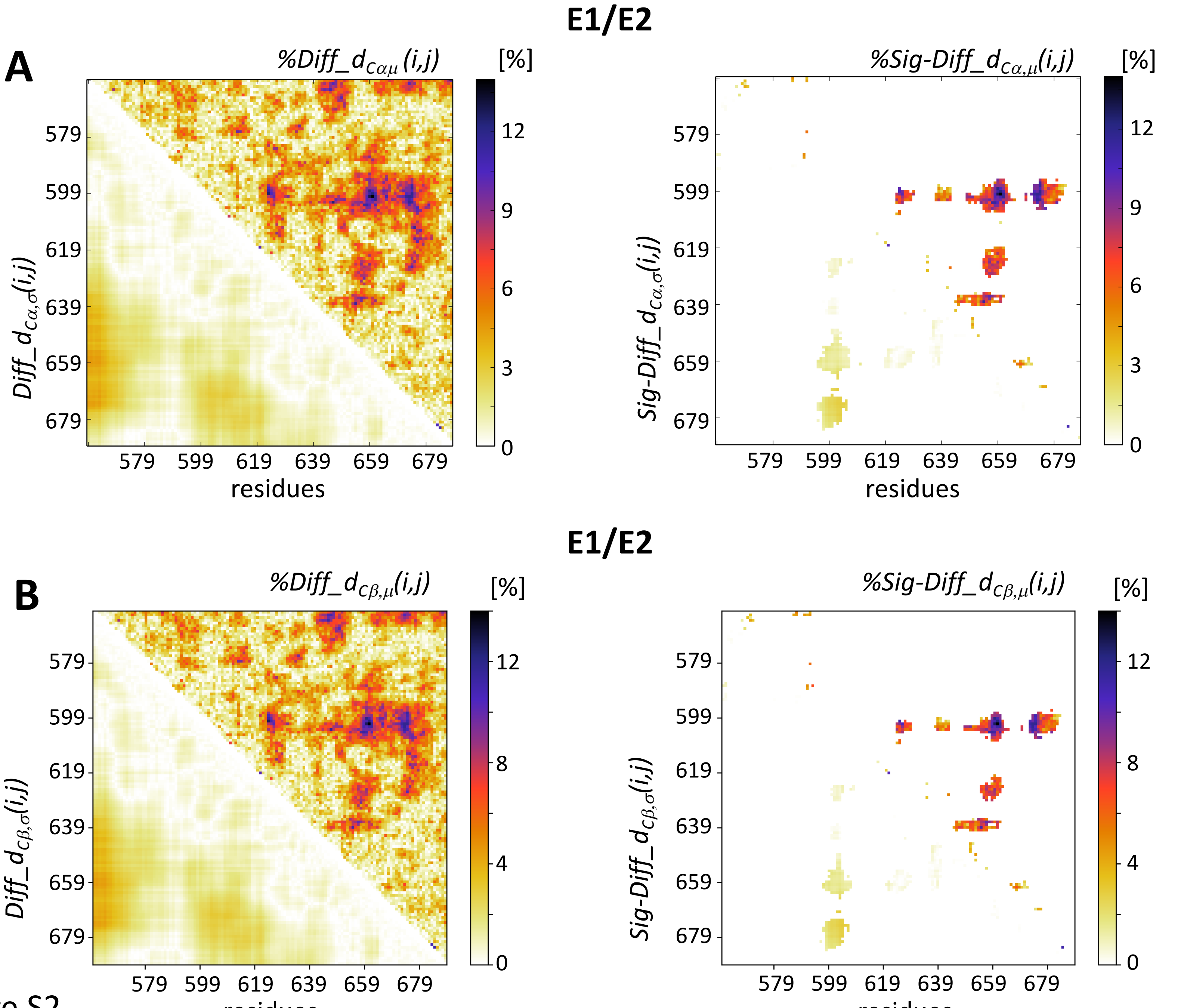
E1/E2
A
%Diff_d_{Cαμ}(i,j)
[%]
Diff_d_{Cα,σ}(i,j)
residues
%Sig-Diff_d_{Cα,μ}(i,j)
Sig-Diff_d_{Cα,σ}(i,j)
E1/E2
B
%Diff_d_{Cβ,μ}(i,j)
Diff_d_{Cβ,σ}(i,j)
%Sig-Diff_d_{Cβ,μ}(i,j)
Sig-Diff_d_{Cβ,σ}(i,j)
579
599
619
639
659
679

Figure S2

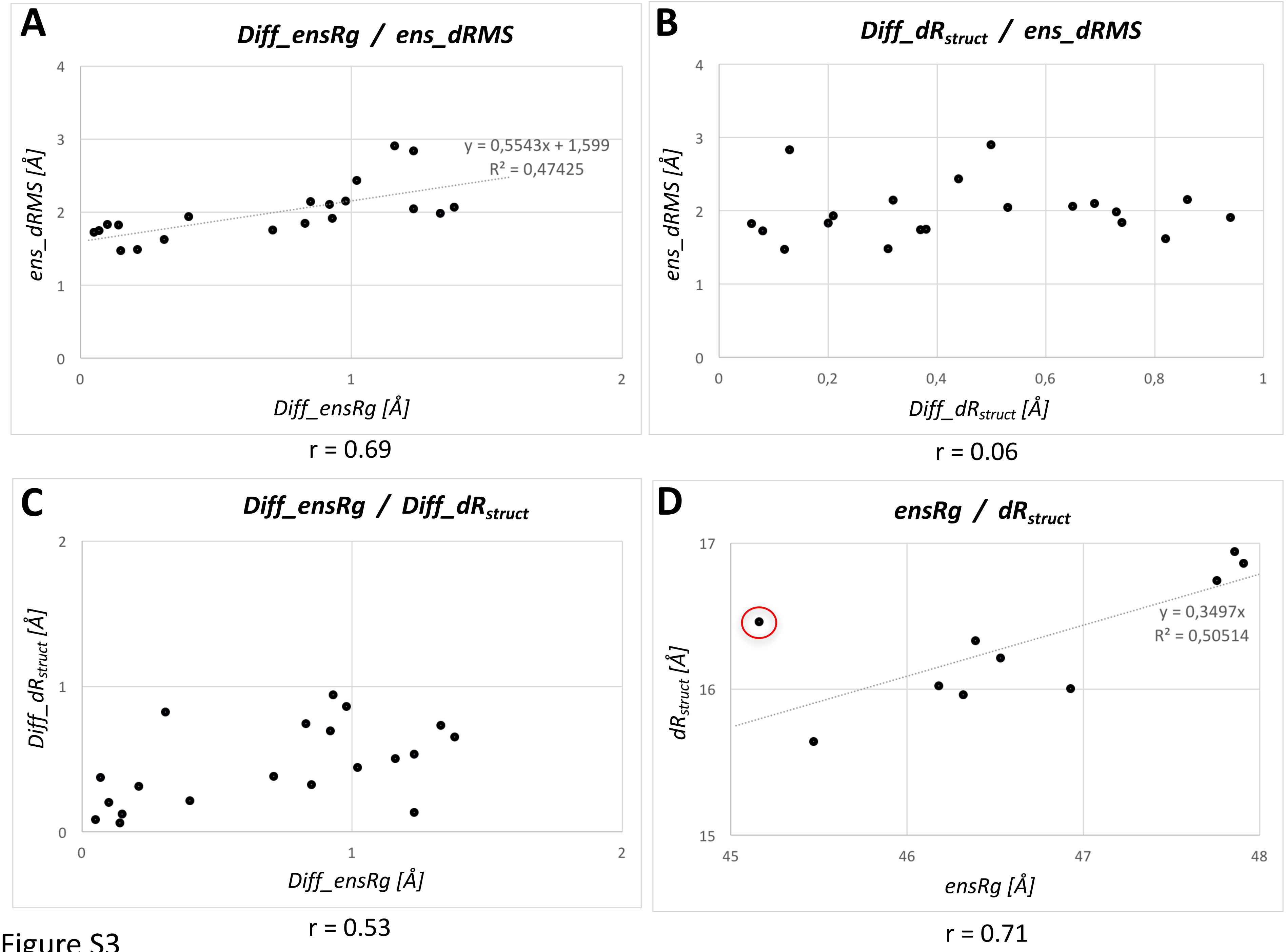
A
Diff_ensRg / ens_dRMS
ens_dRMS [Å]
Diff_ensRg [Å]
y = 0,5543x + 1,599
R² = 0,47425
r = 0.69
B
Diff_dRstruct / ens_dRMS
ens_dRMS [Å]
Diff_dRstruct [Å]
r = 0.06
C
Diff_ensRg / Diff_dRstruct
Diff_dRstruct [Å]
Diff_ensRg [Å]
r = 0.53
D
ensRg / dRstruct
dRstruct [Å]
ensRg [Å]
y = 0,3497x
R² = 0,50514
r = 0.71

Figure S3

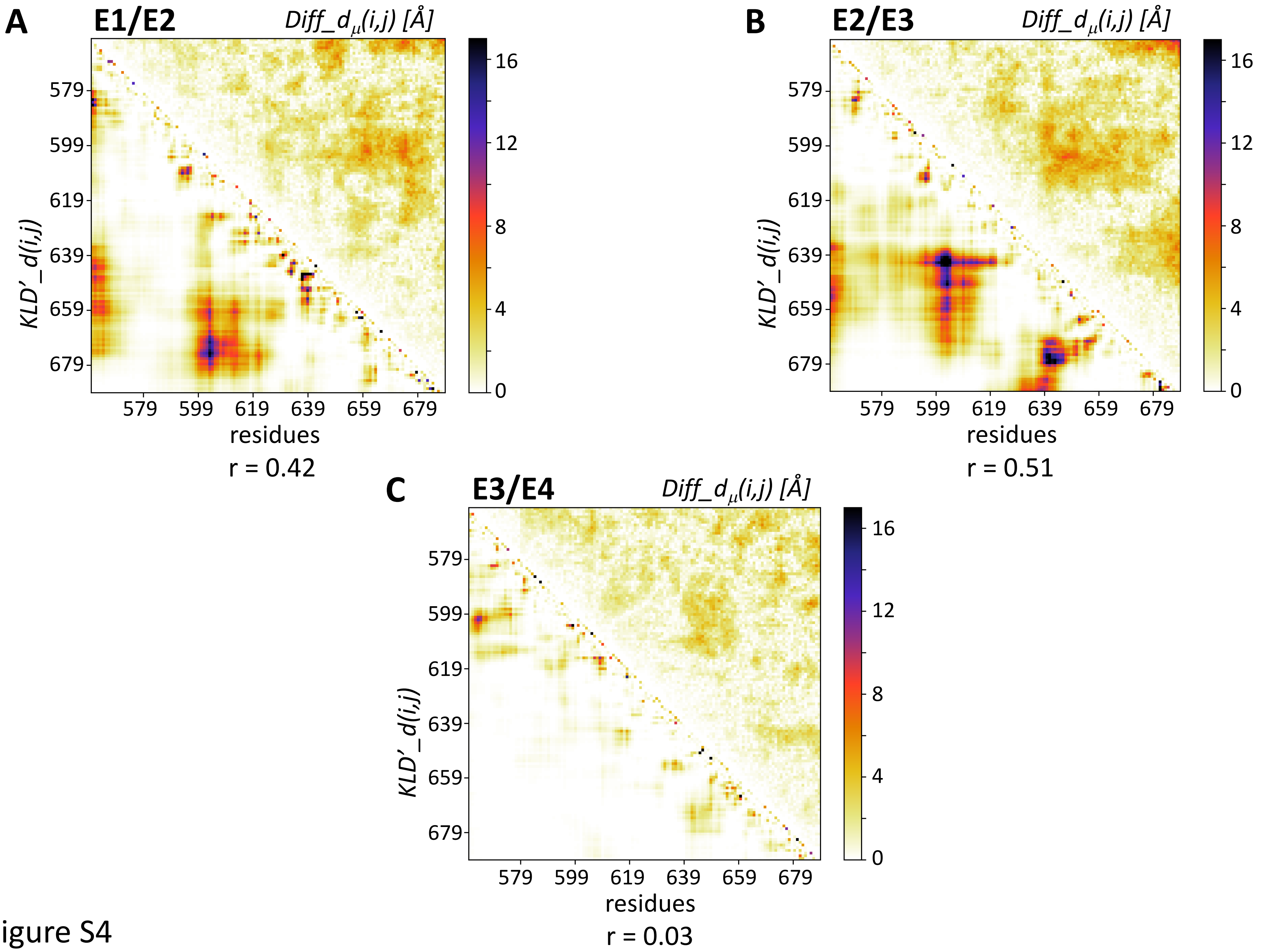
A
E1/E2
Diff_dμ(i,j) [Å]
KLD'_d(i,j)
residues
579 599 619 639 659 679
0 4 8 12 16
r = 0.42
B
E2/E3
Diff_dμ(i,j) [Å]
KLD'_d(i,j)
residues
579 599 619 639 659 679
0 4 8 12 16
r = 0.51
C
E3/E4
Diff_dμ(i,j) [Å]
KLD'_d(i,j)
residues
579 599 619 639 659 679
0 4 8 12 16
r = 0.03

Figure S4

### AA sequences of the MeV N-tail and tau-K18 domain

```
>PED7AAC|NCAP_MEASF Nucleoprotein N-tail OS=Measles virus
MHHHHHHTTEDKISRAVGPRQAQVSFLHGDQSENELPRLGGKEDRRVKQSRGEARESYRET
GPSRASDARAAHLPTGTPLDIDTASESSQDPQDSRRSADALLRLQAMAGISEEQGSDTDTP
IVYNDRNLLD
```

```
>PED6AAC|TAU_HUMAN MAPT tau-K18 segment OS=Homo sapiens
LQTAPVPMPDLKNVKSKIGSTENLKHQPGGGKVQIINKKLDLSNVQSKCGSKDNIKHVPGG
GSVQIVYKPVDLSKVTSKCGSLGNIHHKPGGGQVEVKSEKLDFKDRVQSKIGSLDNITHVP
GGGNKKIE
```

Figure S5

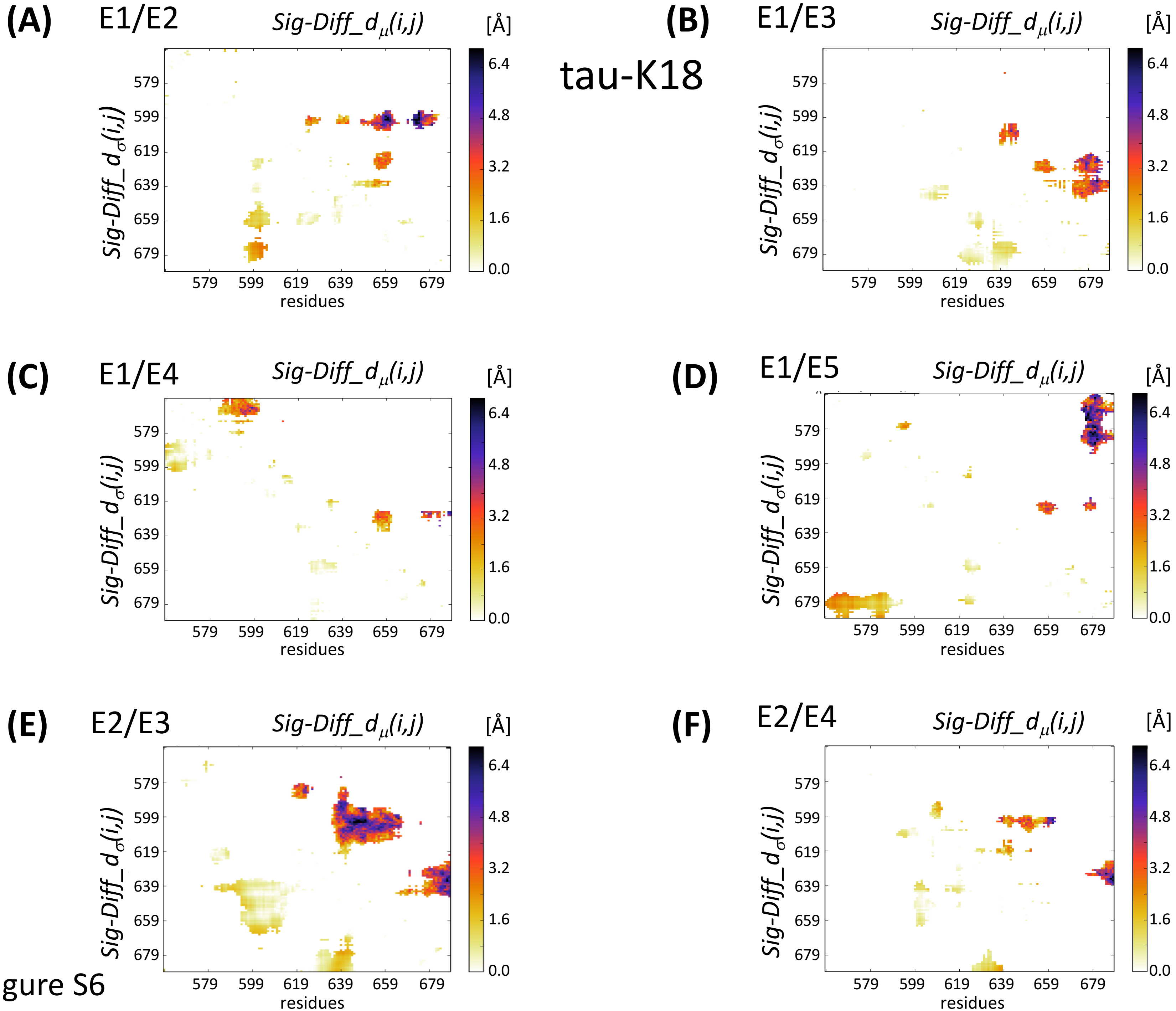
tau-K18
(A) E1/E2
(B) E1/E3
(C) E1/E4
(D) E1/E5
(E) E2/E3
(F) E2/E4
$Sig\text{-}Diff\_d_{\mu}(i,j)$
$Sig\text{-}Diff\_d_{\sigma}(i,j)$
[Å]
6.4
4.8
3.2
1.6
0.0
579
599
619
639
659
679
residues

Figure S6

### tau-K18

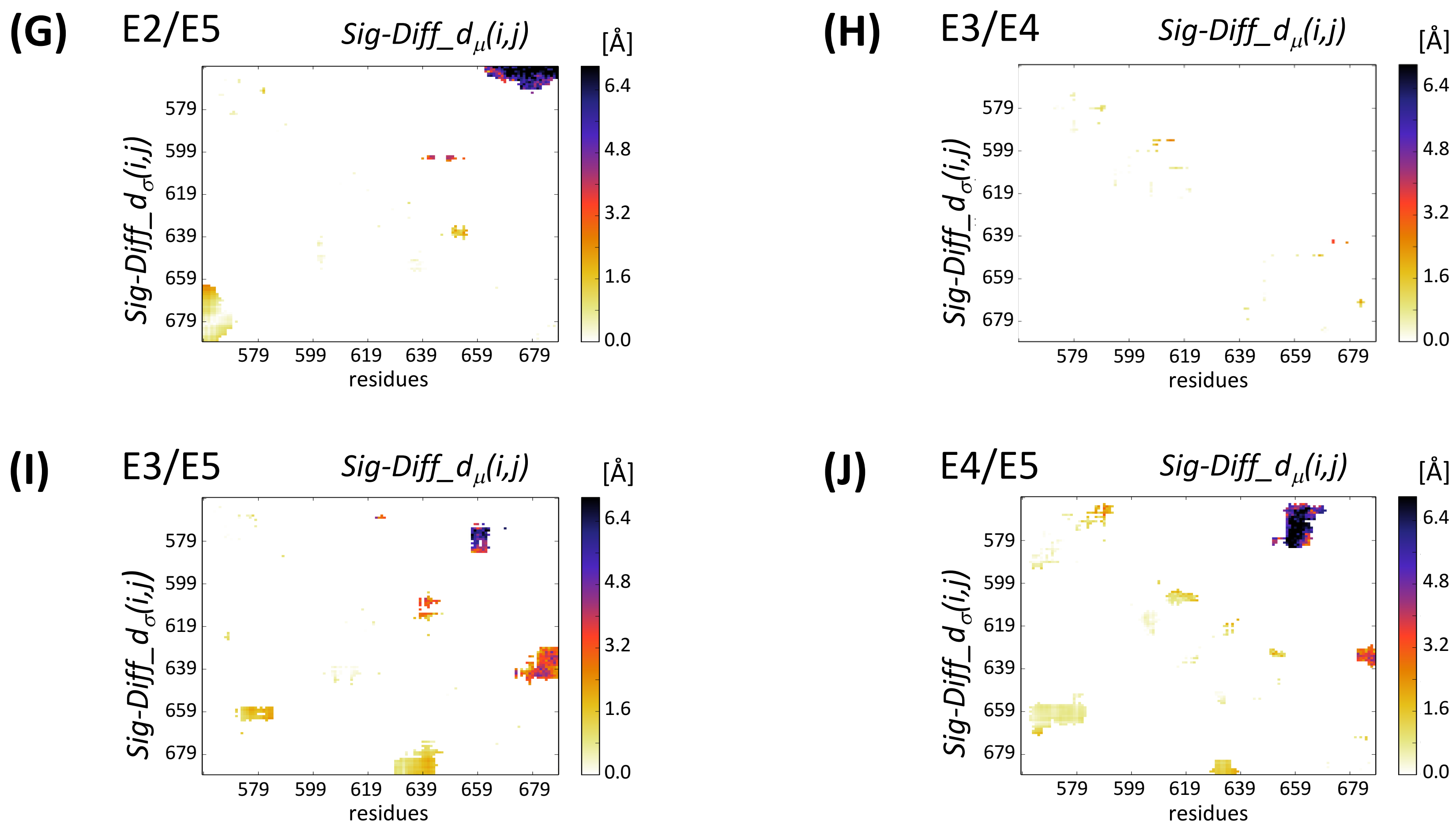


Figure S6

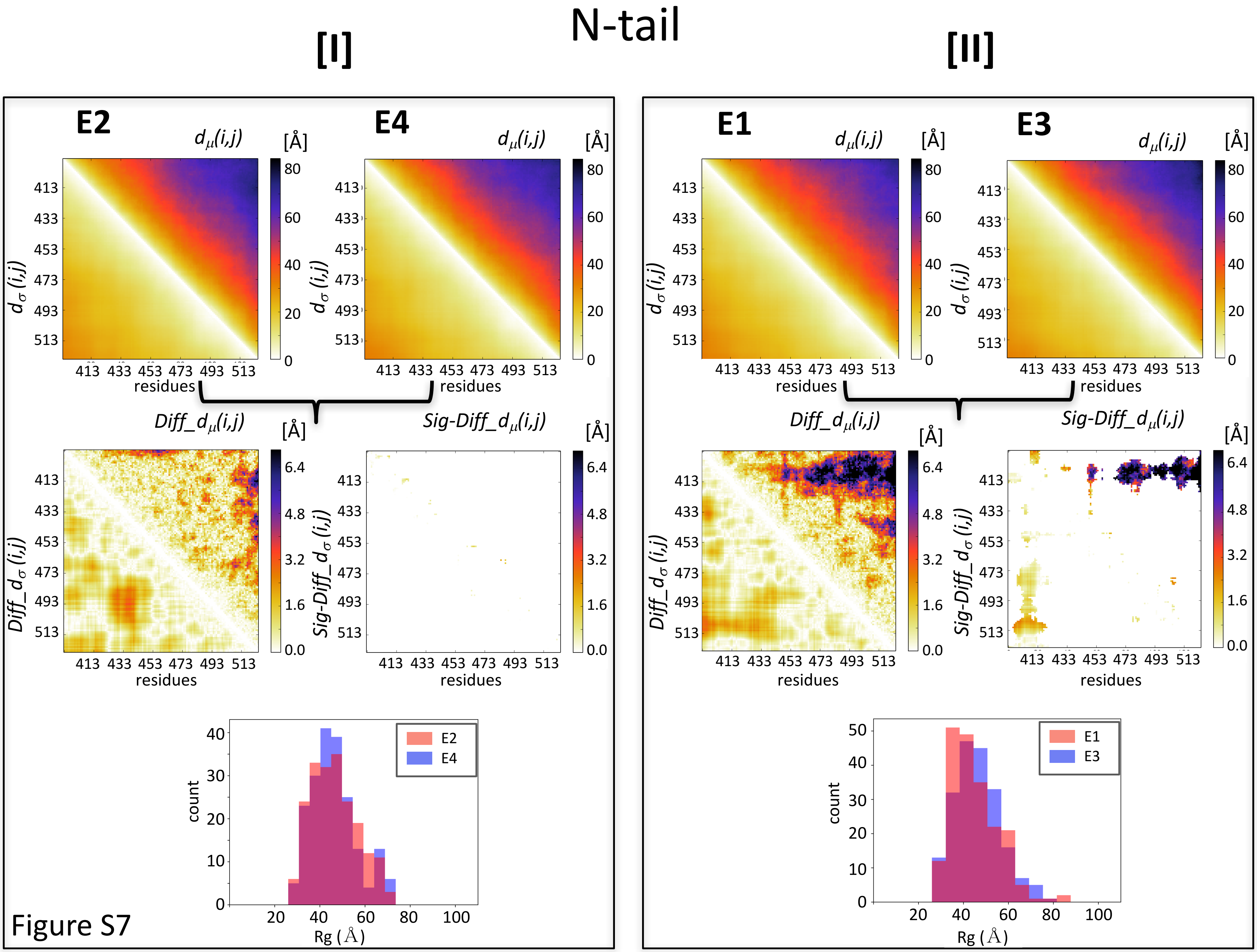
N-tail
[I]
[II]
E2
E4
E1
E3
$d_\mu(i,j)$
$d_\sigma(i,j)$
[Å]
80
60
40
20
0
413 433 453 473 493 513
residues
$Diff\_d_\mu(i,j)$
$Diff\_d_\sigma(i,j)$
$Sig\text{-}Diff\_d_\mu(i,j)$
$Sig\text{-}Diff\_d_\sigma(i,j)$
6.4
4.8
3.2
1.6
0.0
count
Rg (Å)
20 40 60 80 100
E2
E4
E1
E3

Figure S7

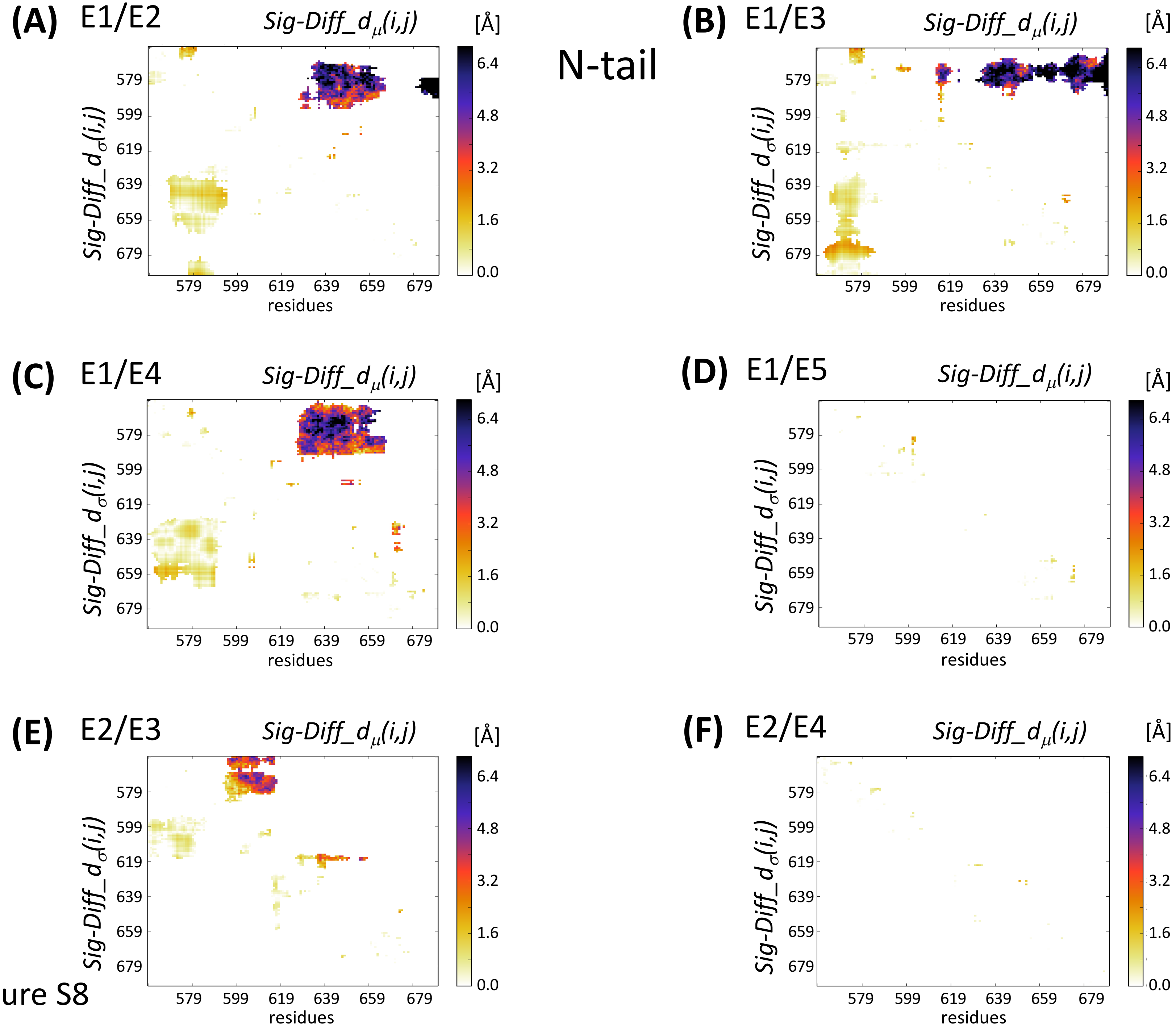

N-tail
(A) E1/E2
(B) E1/E3
(C) E1/E4
(D) E1/E5
(E) E2/E3
(F) E2/E4
Sig-Diff_dμ(i,j)
Sig-Diff_dσ(i,j)
[Å]
6.4
4.8
3.2
1.6
0.0
579
599
619
639
659
679
residues


Figure S8

### N-tail

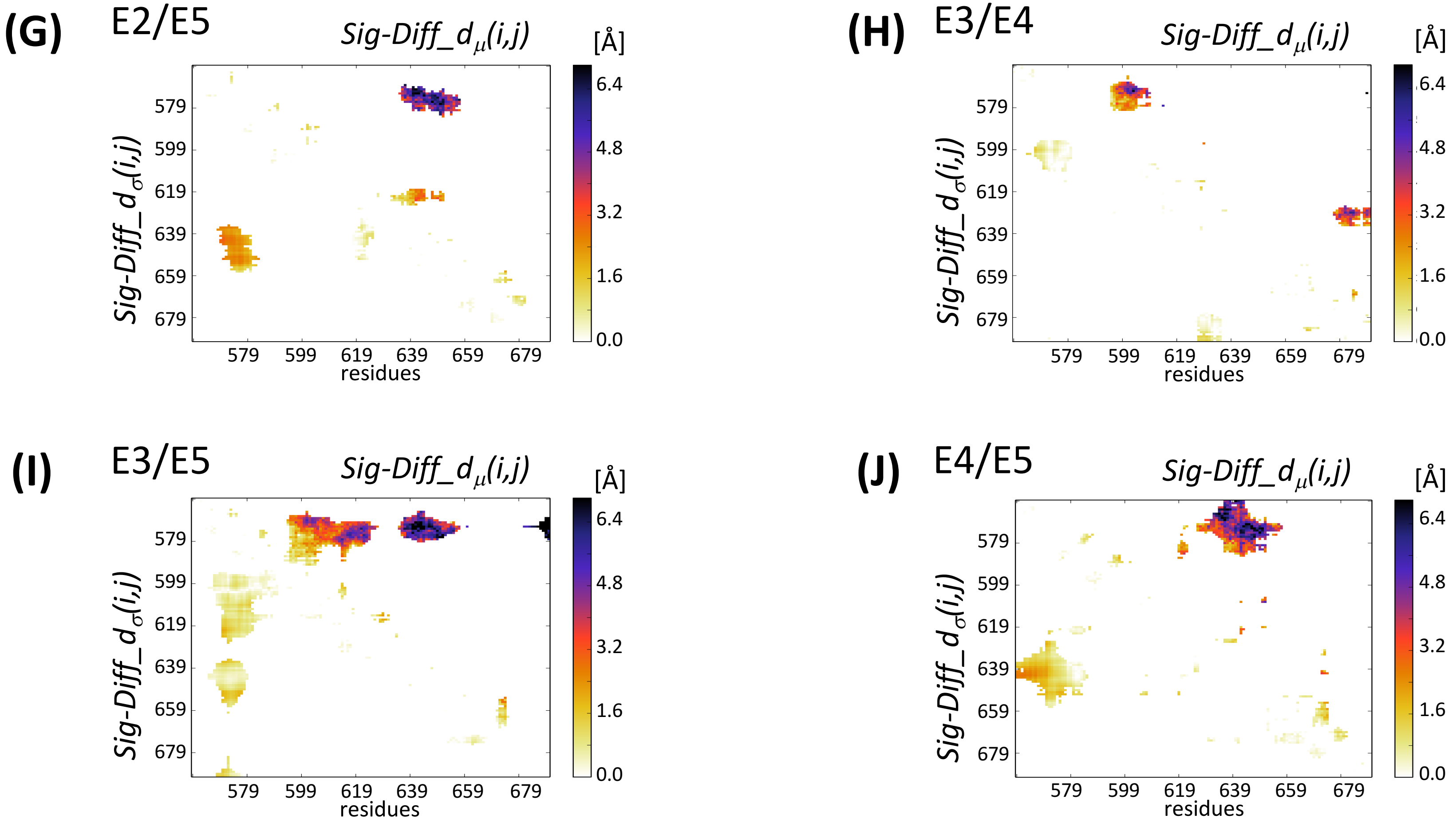


Figure S8

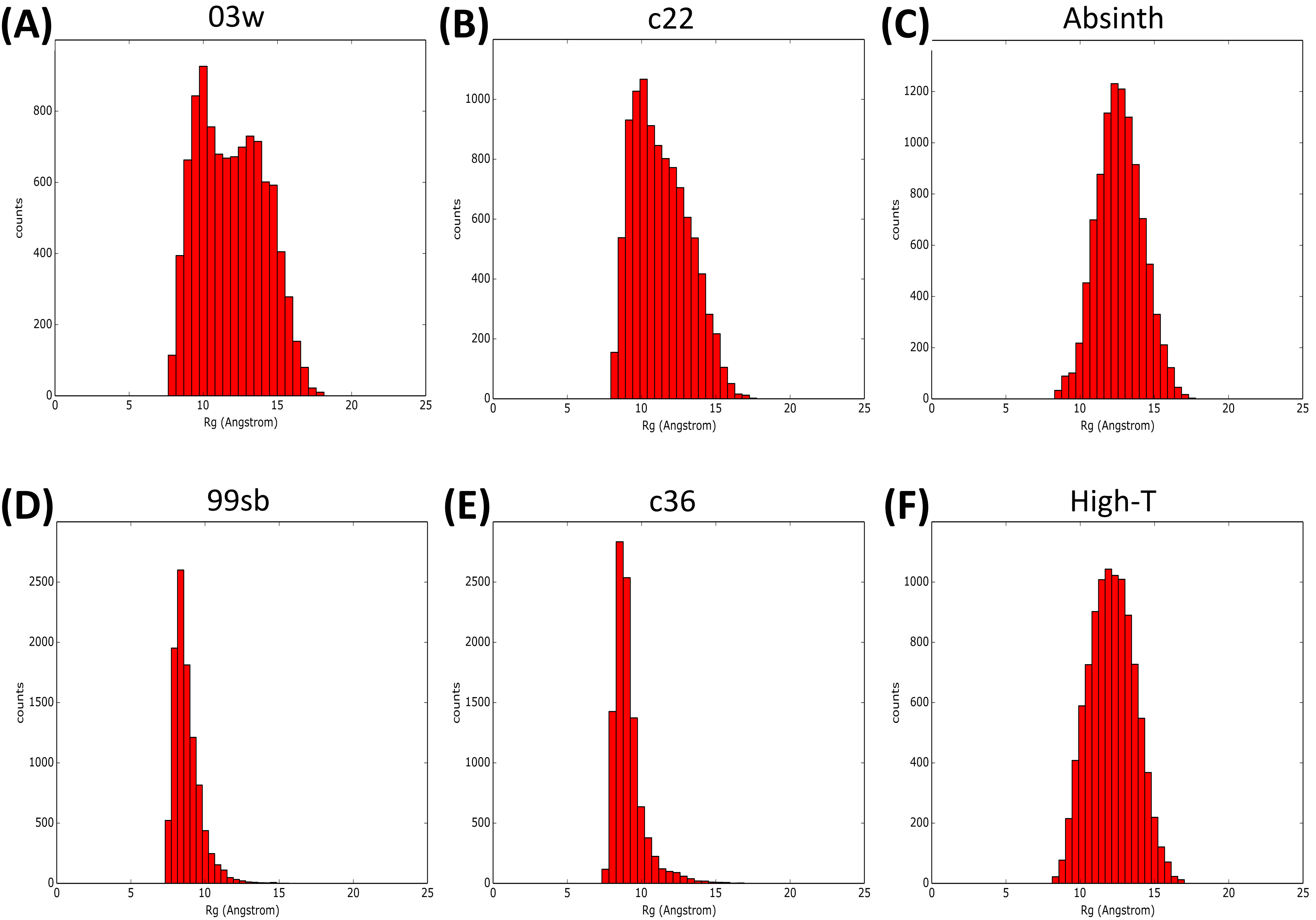


Figure S9

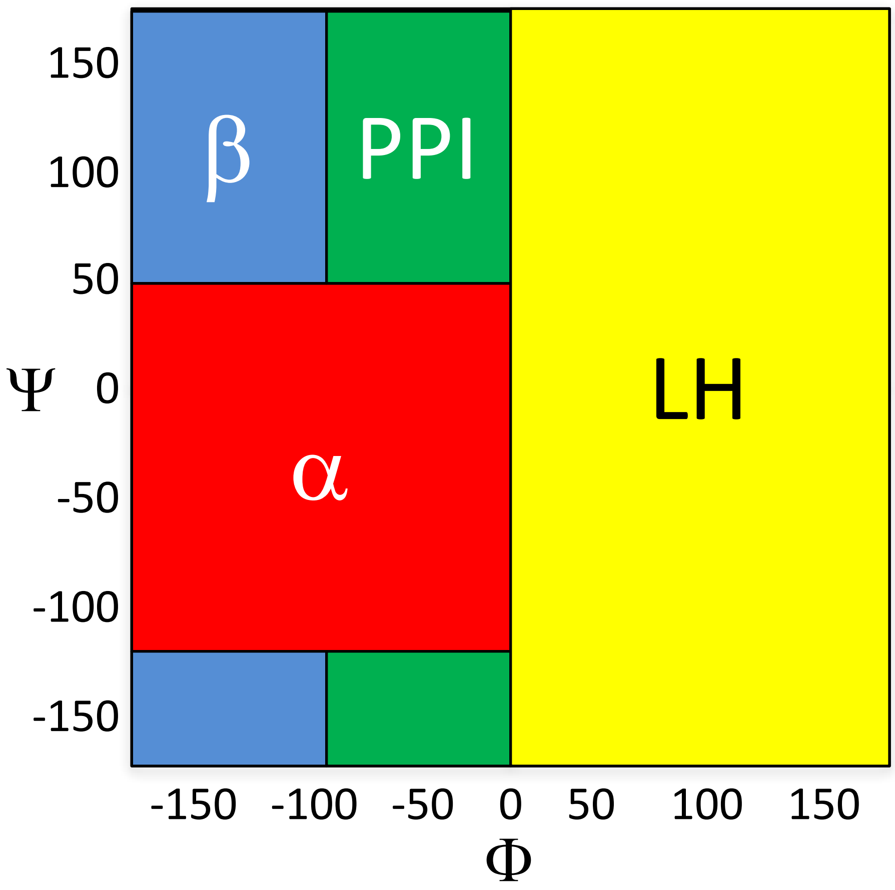


Figure S10
